# A Novel Chronic *in vivo* Oral Cadmium Exposure-Washout Mouse Model for Studying Cadmium Toxicity and Complex Diabetogenic Effects

**DOI:** 10.1101/2022.02.21.481351

**Authors:** Winifred P.S. Wong, Janice C. Wang, Matthew S. Meyers, Nathan J. Wang, Rebecca A. Sponenburg, Norrina B. Allen, Joshua E. Edwards, Malek El Muayed

## Abstract

Type II diabetes mellitus (T2DM) is characterized by insulin resistance, β-cell dysfunction and hyperglycemia. In addition to well known risk factors such as lifestyle and genetic risk score, accumulation of environmental toxicants in organs relevant to glucose metabolism is increasingly recognized as additional risk factors for T2DM. Here, we describe the development of an *in vivo* oral cadmium (Cd) exposure model. It was shown that oral Cd exposure in drinking water followed by washout and high fat diet (HFD) in C57BL/6N mice results in islet Cd bioaccumulation comparable to that found in native human islets while mitigating the anorexic effects of Cd to achieve the same weight gain required to induce insulin resistance as in Cd naïve control mice. Inter individual variation in plasma glucose and insulin levels as well as islet Cd bioaccumulation was observed in both female and male mice. Regression analysis showed an inverse correlation between islet Cd level and plasma insulin following a glucose challenge in males but not in females. This finding highlights the need to account for inter individual target tissue Cd concentrations when interpreting results from *in vivo* Cd exposure models. No effect of Cd on insulin secretion was observed in islets *ex vivo*, highlighting differences between *in vivo* and *ex vivo* cadmium exposure models. In summary, our oral *in vivo* Cd exposure-washout with HFD model resulted in islet Cd bioaccumulation that is relevant in the context of environmental cadmium exposure in humans. Here, we showed that islet Cd bioaccumulation is associated with complex cadmium-mediated changes in glucose clearance and β-cell function. The model described here will serve as a useful tool to further examine the relationship between Cd exposure, islet Cd bioaccumulation, dysglycemia and their underlying mechanisms.

## Introduction

Type II diabetes mellitus (T2DM) is characterized by insulin resistance and dysfunction of insulin-secreting β-cells within pancreatic islets of Langerhans (Petersmann, Muller-Wieland et al. 2019). The development of T2DM is dependent upon the interplay of many factors. Several genetic variations (Todd, Srinivasan et al. 2018, Banerjee, Vats et al. 2019), intrauterine environment (Fernandez-Twinn, Hjort et al. 2019) and lifestyle-related factors including obesity, diet, physical inactivity and smoking have been associated with susceptibility of T2DM (Kolb and Martin 2017). In addition to these risk factors, increasing evidence suggests that exposure to environmental toxicants such as arsenic (Martin, Gonzalez-Horta et al. 2015, Kuo, Moon et al. 2017, Beck, Chandi et al. 2019, Li, Douillet et al. 2020) and cadmium (Cd) (Schwartz, II’yasova et al. 2003) as contributors to the development of T2DM.

To date, studies in humans and animal models describing the relationship between Cd exposure on diabetes are inconsistent. Some epidemiologic studies showed an association between Cd exposure and impaired glucose homeostasis (Bell, Early et al. 1990, Schwartz, II’yasova et al. 2003, Afridi, Kazi et al. 2008, Fitzgerald, Olsen et al. 2020) while others reported the lack of an association (Barregard, Bergstrom et al. 2013, Menke, Guallar et al. 2016). Cadmium is a transition metal with toxic properties that is found at low levels in fruits, grains and vegetables grown on Cd-containing soil (Jolly, Islam et al. 2013), drinking water (Dalmieda and Kruse 2019) and cigarette smoke (Scherer and Barkemeyer 1983). Exposure to environmental cadmium through ingestion of Cd-containing food components and inhalation of cigarette smoke results in gradual accumulation of Cd in various organs including insulin-producing pancreatic islets (Wong, Allen et al. 2017). In mice, short term Cd exposure *in vivo* given as subcutaneous injections resulted in bioaccumulation of Cd in pancreatic islets and dysglycemia (Fitzgerald, Olsen et al. 2020). In dispersed mouse islet cell culture, bioaccumulation of intracellular Cd was associated with a blunted glucose-stimulated insulin secretion (El Muayed, Raja et al. 2012). Further studies examining the effects and mechanisms of sustained, oral Cd in insulin-producing islets at environmentally relevant concentrations while avoiding acute systemic toxicity are warranted. Cadmium has a long tissue half-life of up to 30 years and thus, results in gradual Cd accumulation in human tissues over decades (Elinder, Lind et al. 1976, Jarup and Akesson 2009) with the highest levels in the kidney cortex reached at age 50 (Friberg 1984). Replicating similar rates of tissue Cd accumulation in a murine model is of high importance for examining the effect of Cd at tissue levels but at the same time, poses unique challenges relating to the long half-life of Cd and its low oral bioavailability of 2-4% (Schilderman, Moonen et al. 1997).

In the present study, we established a novel *in vivo* oral Cd exposure murine model with an initial oral Cd exposure phase followed by a washout phase with high fat feeding aimed at replicating chronic islet Cd accumulation in the setting of insulin resistance in humans over a reasonable time frame. Our Cd exposure-washout with high fat feeding mouse model results in near baseline systemic Cd levels while achieving islet Cd bioaccumulation that are within range of those found in native human islets from the general population (Wong, Allen et al. 2017). We used this model to investigate the effects of Cd bioaccumulation in pancreatic islets on β-cell function at islet Cd concentrations that are relevant to human environmental Cd exposure.

## Material and methods

### Cadmium exposure in mice

5-6 weeks old, C57BL/6N mice were exposed to either vehicle or 0.5mM (=56.2ppm Cd, 56.2 mg/L Cd) or 1mM CdCl_2_ (112.4ppm Cd, 112.4 mg/L Cd) (Rigaku, Cat #: 1008154) in deionized drinking water (18.2µΩ, Millipore) acidified with HCl (30µM, Fisher Scientific) for varying duration as indicated. During Cd exposure, mice were fed regular chow (Envigo, Cat #: 7912) *ad libitum* unless otherwise stated. Experiments were carried out in several successive cohorts designed to test the effect of various exposure parameters as detailed in Table 1. In some experiments, mice were transitioned from CdCl_2_-containing drinking water to vehicle drinking water during a subsequent washout phase. Mice were fed either regular chow (Envigo, Cat #: 7912) or adjusted calories high fat rodent chow (Envigo, Cat #: TD88137, TD180240, 42% from fat) *ad libitum* during the exposure and/or washout phase as described. Drinking water was changed once a week to ensure freshness throughout all experimental phases. Metal analysis of regular and high fat chow used are summarized in Suppl. Table 3. At the conclusion of the study, mice were euthanized and tissues were collected for further analysis. All animal experiments were reviewed and approved by the Institutional Animal Care and Use Committee of Northwestern University.

**Table 1:**
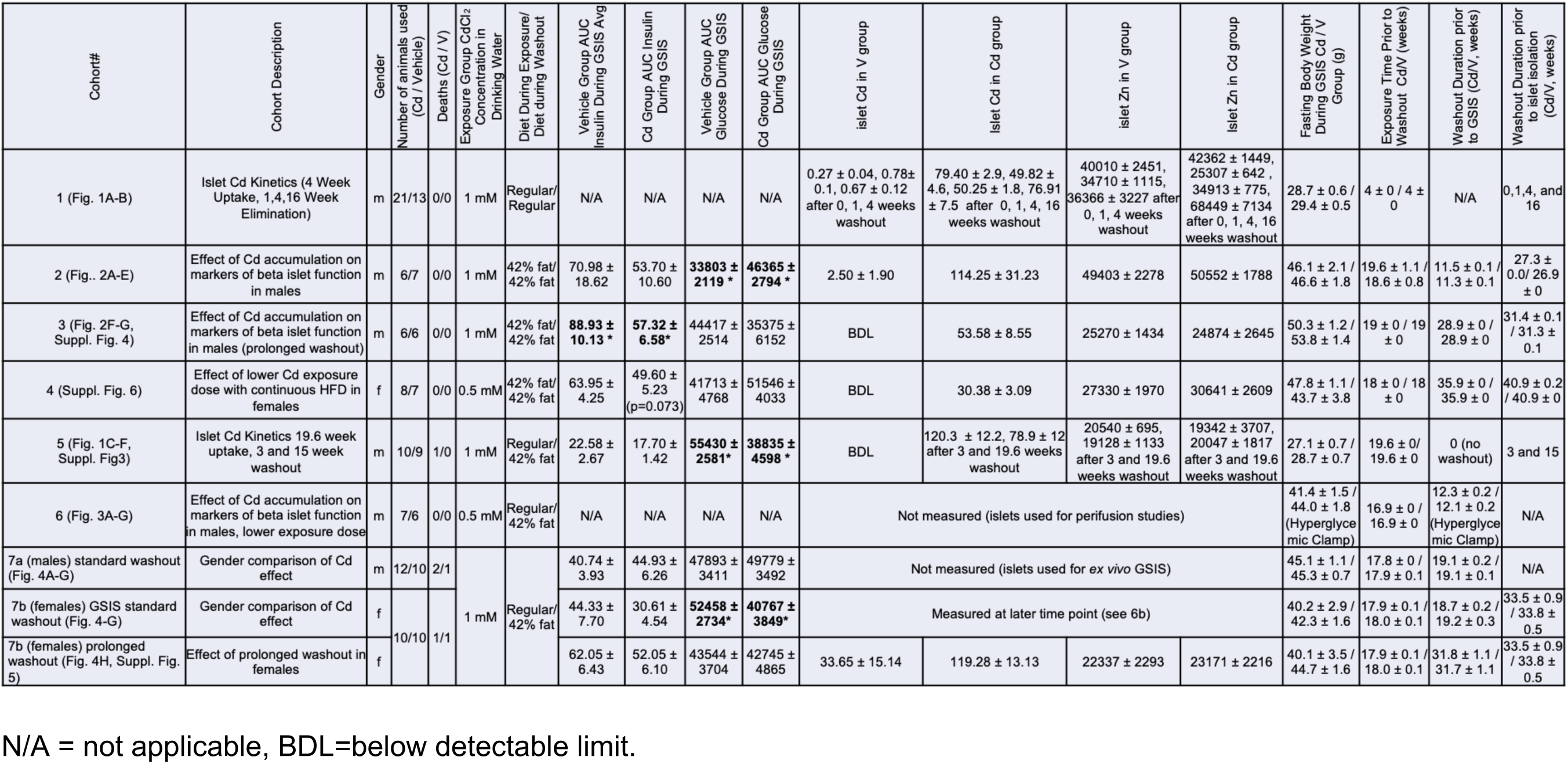
List of cohorts with their relevant experimental and islet outcome parameters. Data are expressed as mean ± SEM. *p<0.05.

### *In vivo* glucose tolerance tests and glucose-stimulated insulin secretion

Standard glucose tolerance test (GTT) was performed by injecting a glucose bolus (50% dextrose, pH 4.2, Hospira, Cat #: 000622) into the intraperitoneal cavity at a dose of 1g/kg body weight following an overnight fast of 16-18 hours (Ayala, Samuel et al. 2010). Blood was drawn through a tail capillary before glucose injection (t=0 min) and at 2, 5, 15, 30, 60 and 120 minutes post-glucose load for blood glucose determination. *In vivo* glucose-stimulated insulin secretion (GSIS) was performed using a higher glucose dose of 3g/kg body weight (Ayala, Samuel et al. 2010). Blood sampling time points from tail capillary for glucose measurements during GSIS were similar to that of GTT tests. Blood glucose concentrations were measured using an Accu-Chek Nano glucose meter (Roche Diagnostics). For GSIS, 20μl of blood was also collected and separated in heparinized microvette tubes (Sarstedt, Germany, Cat #: CB300LH) before glucose injection (t=0min) and at 2, 5, 15 and 30 minutes post-glucose load for plasma insulin analysis. Plasma samples were stored at −80°C until measurements of plasma insulin that was performed using ELISA kits (Crystal Chem, Cat #: 90080). In some experiments, additional plasma samples were obtained for determination of proinsulin levels (Mercodia, Cat #: 10-1232-01).

### Hyperglycemic clamp

Hyperglycemic clamps were performed by the Mouse Metabolic Phenotyping Center (MMPC), Vanderbilt University, Tennessee. Briefly, 5 days prior to clamp studies, catheters were implanted into the carotid artery and jugular veins for sampling and infusion respectively. Hyperglycemic clamps were performed in unrestrained, conscious mice following 5 hours fast (Ayala, Samuel et al. 2010). At t=0 min, blood glucose was raised and continuous glucose infusion rate adjusted to maintain hyperglycemic plateau between 250-300mg/dl (13.8-16.7mM) based on arterial blood glucose measurements at 10 min intervals. Mice received heparinized saline-washed mouse erythrocytes from donors at 5μl/min to prevent a drop in hematocrit. Baseline arterial blood glucose, insulin and c-peptide were measured at −15 and −5 minutes prior to clamp. Arterial blood sampling for clamp insulin was measured at 5, 10, 15, 20, 40, 60, 80, 100 and 120 minutes during clamp. Arterial c-peptide was measured at 15, 100 and 120 minutes during clamp. Plasma insulin and plasma c-peptide were measured using radioimmunoassay (EMD Sigma, Cat #: RI-13K) and bead immunoassay (Luminex, R&D Systems) respectively that were used according to manufacturer’s instructions. Following completion of clamp studies, mice were returned to regular housing for at least 48 hours with food and water available *ad libitum* prior to islet perifusion studies.

### Pancreatic islet isolation

Islet isolation was performed as previously described (El-Muayed, Raja et al. 2012). In brief, C57BL/6N mice were anaesthetized with intraperitoneal injection of ketamine (10mg/ml)/xylazine (1mg/ml) and the abdominal cavity opened to expose the common bile duct. Pancreas was perfused with 3ml collagenase (0.5mg/ml, Sigma, Cat #: C7657) dissolved in HBSS buffer (Corning, Cat #: 21-020-CV) with phenol red through the common bile duct. Inflated pancreases were excised, washed in phenol red-free HBSS buffer (Gibco, Cat #: 14175-103) and digested at 37°C for 15 minutes. Islets were purified from digested pancreas tissue by centrifugation in Biocoll density gradient (EMD Millipore, Cat #: L6155) at 4°C, 2500rpm for 20 minutes. Purified islets were hand-picked under a microscope (Leica Microsystems) and transferred into clean microcentrifuge tubes. Islets were washed 3 times in ice-cold PBS (Corning, Cat #: 21-031-CV) at 4°C, 500xg for 2 minutes and stored at −20°C for ICP-MS metal analysis or used for GSIS experiments as described.

### Islet perifusion

Islet perifusion was performed by MMPC Core Facility, Vanderbilt University, Tennessee. In brief, following isolation, islets were handpicked under the microscope for functional assessment. Islet function was assessed by perifusion using approximately 50 islets (∼55 islet equivalents (IEQ)/per perifusion chamber) at a baseline of 5.6mM glucose, and stimulated with 16.7mM glucose, 16.7mM glucose + 100μM 3-isobutyl-1-methyl-xanthine (IBMX, Sigma, Cat #: I5879), 16.7mM glucose + 20mM Arginine/HCl (Sigma, Cat #: A6969), 5.6mM glucose + 300μM Tolbutamide (Sigma, Cat #: T0891), and 5.6mM glucose + 20mM KCl. Perifusate fractions were collected at the flow rate of 3ml/min and insulin was measured using radioimmunoassay kit (Millipore, Cat #: RI-13K). Insulin secretion were normalized to IEQ.

### Measurement of islet and blood metal content

Measurements of islet Cd were performed as previously reported (Wong, Allen et al. 2017). In brief, 100 islets or 30μl of blood was collected in metal-free, trace metal grade, HCl-washed, 1.7ml polypropylene tubes (Denville, Cat #: C-2170). Ceramic scissors and plastic instruments were used for tail nicking and blood collection. Blood and islet samples were dried and heated at 80°C for 30 minutes. Dried blood and islet samples were dissolved in ultra trace elemental grade (TEG) 68% HNO_3_ (Fisher Scientific, Cat #: A467-500), heated to 80°C for 30 minutes followed by dilution with trace metal grade water (EMD, Cat #: WX0003-6) and stored at −20^°^C. Samples were further diluted with trace metal grade water acidified with TEG HNO_3_ to a final concentration of 2.765N prior to submission for Inductively Coupled Plasma Mass Spectrometry (ICP-MS) analysis, as previously reported (Wong, Allen et al. 2017). Islet metal concentrations were normalized to total islet protein. Total islet protein in hydrolyzed samples was measured using a BCA assay kit (Thermo Scientific, Cat #: 23235) following pH neutralization with NaOH. Accuracy of BCA assay protein measurements using this procedure was validated using BSA standards. Negative ICP-MS output values were imputed to zero. Certified metal-free pipette tips (VWR, Cat #: 53508-918), conical tubes (VWR Cat #: 89049-170, 89049-174) and 1.7ml polypropylene microcentrifuge tubes (Denville, Cat #: C-2170) were used throughout the collection and analysis process. All steps were performed in an epoxy-coated laminar flow hood with trace element grade air filter (Baker, UK).

### ^67^Zn turnover in MIN6 cells

MIN6 cells, an immortalized murine β-cell line (ATCC, Menassas, VA) (Miyazaki, Araki et al. 1990) were exposed to either vehicle or 0.5µM CdCl_2_ for 48 hours as previously described (El Muayed, Raja et al. 2012). Growth media was supplemented with 10μM ZnCl_2_ to achieve a total Zn^2+^ concentration of 15μM in culture (native media contains 5μM Zn^2+^). This exposure regimen results in significant accumulation of Cd (El Muayed, Raja et al. 2012). Cells were harvested 48 hours after the start of Cd exposure. At 1, 2 or 4 hours prior to harvest, culture media was changed and fresh media containing the rare, non-radioactive ^67^Zn tracer was added as ^67^ZnCl_2_ at a concentration of 10μM (in addition to the 5μM natural Zn mix found in media = 66.6% ^67^Zn). At harvest, samples were washed twice in PBS and incubated in 0.05% trypsin containing 0.5mM EDTA (37°C, 4 min, Gibco Cat #: 25300054). Following trypsinization, samples were transferred into fresh microcentrifuges tubes, centrifuged (500 RCF, 3 min) and further washed twice with PBS. Pellets were stored at −20°C until processed for metal profile including Zn isotopes by ICP-MS. The rate of divalent ^67^Zn uptake was used as a proxy measure for Zn turnover.

### Statistical analysis

Area Under the Curve (AUC) for glucose and insulin curves during GSIS were calculated using the trapezoid rule. Two-tailed t-tests were performed to compare differences between control and test groups (Prism, v9.3.1). One way ANOVA with post hoc Tukey’s test for multiple comparison was performed to compare differences between more than two groups. Where appropriate, two-way ANOVA with repeated measures was performed to compare glucose and insulin secretion over time (Prism, v9.3.1). Linear regression analysis was performed between islet Cd levels, islet Zn levels, glucose AUC and insulin AUC in both male and female datasets (SAS). Values are expressed as mean ± standard error of the mean (SEM). p<0.05 were considered statistically significant.

## Results

### Development of the newly established *in vivo* Cd exposure-washout model

In an initial pilot experiment, we exposed male C57BL/6N mice to either vehicle (V) or 1mM CdCl_2_ in drinking water continuously for 12 months while being fed normal chow. There were no differences in either glucose tolerance, glucose area under the curve or body weight between Cd-exposed and V controls in this initial model of Cd exposure without diet-induced insulin resistance (Suppl. Fig 1A-C, not included in Table 1). Next, we performed experiments aimed at examining the effects of Cd exposure with high fat diet-induced insulin resistance (Ayala, Samuel et al. 2010). To ameliorate the well described anorexigenic effects of Cd, we employed an initial Cd exposure phase, during which mice were exposed to either 0.5mM or 1mM CdCl_2_ in their drinking water to induce Cd tissue accumulation to environmentally relevant levels, followed by a washout phase (without Cd exposure). The purpose of the washout phase is to mitigate the anorexigenic and other systemic effects of Cd at the time of phenotyping. This is a metabolic study and body weight is an important parameter that affects and/or influences metabolic outcomes. Initial cohorts employed high fat diet (HFD, 42% fat) throughout the exposure and washout phases (Table 1, Cohorts 2-4). Later cohorts employed regular chow during the exposure phase and transitioned to HFD only during the washout phase in order to induce weight gain closer to the timepoint of phenotyping (Table 1, Cohorts 5-7). Animal cohorts and their experimental parameters that were performed in this study are summarized in Table 1. All diets employed in the current study were analyzed and found to be low in Cd content (Suppl. Table 3). The high fat diet had lower Zn concentration compared to the regular chow.

### *In vivo* islet and blood Cd kinetics

Initial experimental cohorts were aimed at establishing the *in vivo* kinetics of Cd in blood and pancreatic islets. To do this, we exposed male mice to either V or 1mM CdCl_2_ for 4 weeks followed by a washout phase (without High Fat Diet (HFD)) and measured Cd levels in blood and pancreatic islets over the course of both the exposure and washout phases to establish islet Cd accumulation and elimination kinetics (Table 1, Cohort #1). Following 4 weeks of Cd exposure, we observed a significant increase in both islet Cd accumulation (79.40 ± 3.40nmol/g protein vs 0.27 ± 0.05nmol/g protein, p<0.001, Fig 1A) and systemic blood Cd (0.9434 ± 0.0078μM vs 0.0033 ± 0.0003μM, p<0.001, Fig 1B) in Cd-exposed mice compared to V controls. However, at 16 weeks into the washout phase (without HFD), we continue to observe an increase in islet Cd accumulation (Fig 1A) but with systemic blood Cd levels (Fig 1B) returning to near baseline levels in Cd-exposed mice. Overall, this demonstrates that the bioaccumulated Cd in murine islets as a consequence of oral Cd exposure *in vivo* remains and persists in islets long after the cessation of Cd exposure despite the return of systemic Cd levels to baseline. For comparison, Cd concentration median and range found in native human islets are 26.1nmol/g protein and 4.4-94.7nmol/g protein respectively (Wong, Allen et al. 2017). Therefore, islet Cd concentrations are comparable but lower than the high end of concentrations found in human islets from the general US population (Wong, Allen et al. 2017). Aiming to achieve islet Cd concentration towards the upper end of those found in human islets, we exposed a separate cohort of male mice with either V or 1mM CdCl_2_ in their drinking water for 19.5 weeks followed by a washout phase (Table 1, Cohort #5). HFD was introduced in this cohort to assess Cd kinetics and weight trajectory under diet-indued hypercaloric conditions. We euthanized two subgroups of mice at 3 and 15 weeks into the washout phase. At 3 weeks into the washout phase, both islet Cd (p<0.001, Fig 1C) and blood Cd levels (Fig 1D) were elevated in Cd-HFD mice compared to V-HFD controls. At 15 weeks into the washout phase, islet Cd levels remain significantly elevated in Cd-HFD mice (p<0.01, Fig 1C) while systemic Cd levels returned to near baseline values (p<0.05, Fig 1D) when compared to V-HFD controls. Islet Cd bioaccumulation in Cd-HFD mice at both 3 and 15 weeks into the washout phase were observed in the absence of significant differences in islet content of Zn, an essential metal that shares many of the transport and buffering pathways with Cd (Fig 1E). Body weight gain were similar between V-HFD and Cd-HFD groups throughout the exposure and washout phases with the exception of one time point towards the end of the Cd exposure phase (Fig 1F, p<0.05, 33.5 ± 2.4 g vs 29.8 ± 2.6 g for V-HFD and Cd-HFD groups respectively). As a consequence, we determined that the best time to introduce the washout phase is just prior to the development of the Cd-induced weight loss observed in Cd-exposed mice i.e. between 17-19 weeks of Cd and to restrict the HFD feeding to the washout phase while using regular chow during the Cd exposure phase. As expected in the setting of lower body weight at the end of the exposure period, Cd-exposed mice showed a lower glucose curve during glucose-stimulated insulin secretion (GSIS) at the end of the Cd exposure phase compared to V control mice (Suppl. Fig 3A-E).

**Fig. 1:**
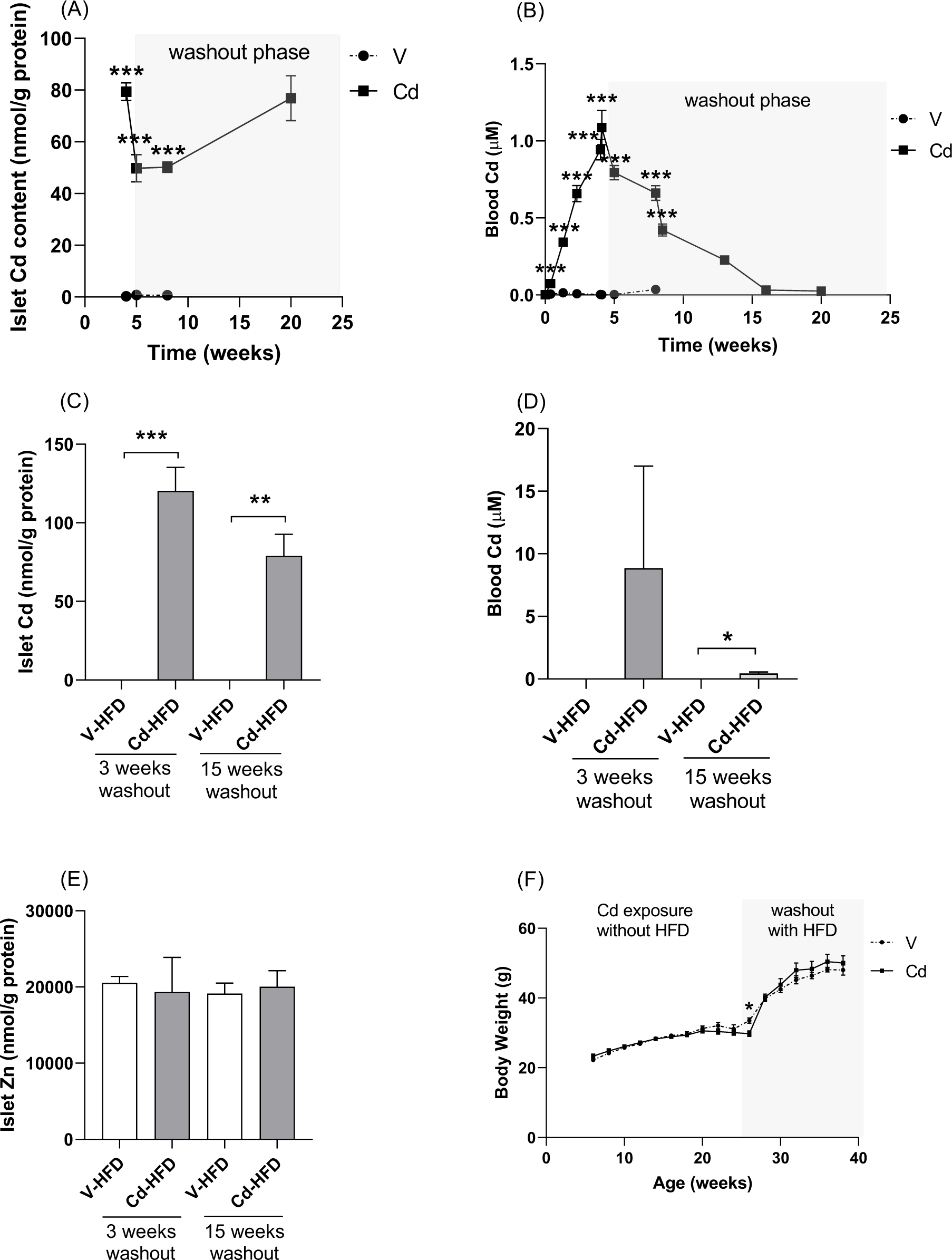
(A) Islet Cd content (normalized to total islet protein) and (B) blood Cd levels in male mice exposed to either V (n=4-5) or 1mM CdCl_2_ (n=4-8) for 4 weeks followed by 16 weeks of washout (Table 1, Cohort #1). (C) Islet Cd content, (D) blood Cd levels, (E) islet Zn content and (F) body weight curve in male mice exposed to either V (n=3-6) or 1mM CdCl_2_ (n=3-6) for 19.5 weeks on regular chow followed by 3 and 15 weeks washout with high fat diet (HFD, Table 1, Cohort #5). V-HFD, Vehicle-exposed followed by washout and HFD; Cd-HFD, Cd-exposed followed by washout and HFD. Data are expressed as mean ± SEM. *p<0.05, **p<0.01, ***p<0.001.

### Cadmium results in transient impairment in glucose clearance in the setting of insulin resistance

To examine the effects of islet Cd accumulation in the setting of insulin resistance in our exposure-washout model, we exposed male mice to either V (n=7) or 1mM CdCl_2_ (n=6) in their drinking water simultaneously with HFD. Mice were exposed to V or Cd for 19.1 ± 0.7 weeks followed by a washout phase (Table 1, Cohort #2). During the washout phase, Cd-exposed mice were placed on vehicle drinking water and both mice groups continue to receive HFD. To characterize the metabolic parameters during the washout phase, we performed metabolic studies in the same cohort of mice at two washout time points (about 12 and 24 weeks into the washout phase). At 11.4 ± 0.1 weeks into the washout phase with HFD, we performed GSIS (3g/kg ip glucose challenge) in both the V and Cd-exposed mice. Cd-HFD mice showed higher glucose values (p<0.05, Fig. 2A, p<0.01 Fig. 2B) without changes in either plasma insulin (Fig. 2C) or fasting body weight (Fig. 2D). However, at 24 weeks into the washout phase with HFD, Cd-HFD mice did not show differences in glucose levels following a glucose tolerance test (Fig. 2E, 1g/kg ip glucose challenge, insulin not measured, per standard glucose tolerance test protocol (Ayala, Samuel et al. 2010). Another cohort of male mice underwent the same procedure with 1mM CdCl_2_ exposure for 19 weeks followed by GSIS after a longer washout period of 28.9 ± 0 weeks and islet isolation at 31.3 ± 0.1 weeks (Table 1, Cohort #3). As with our initial experiments (Fig. 1C), islet Cd (53.6 ± 9.4nmol/g protein, p<0.001, Fig. 2F) in Cd-HFD mice remained elevated compared to V-HFD controls at 31 weeks into the washout phase despite systemic Cd levels being close to baseline (−0.0316 ± 0.0004μM vs. 0.06 ± 0.03μM, p<0.01, Fig. 2G). A significantly lower plasma insulin level (Suppl. Fig 4B) during GSIS was observed in Cd-HFD mice (insulin AUC 57.3 ± 6.6 ng/mL*min) compared to V-HFD controls (88.9 ± 10.1ng/mL*min) (Suppl. Fig 4D, p<0.05) without changes in either plasma glucose levels (Suppl. Fig 4A) or fasting body weight (Suppl. Fig 4E). We exposed a subsequent female cohort to a lower CdCl_2_ concentration of 0.5 mM in drinking water for 18 weeks followed by a washout period with HFD feeding for 35.9 weeks (Table 1, Cohort #4). The lower exposure dose was chosen to account for the higher bioavailability of Cd in females. Here, a significantly lower plasma insulin value at 15 minutes (Suppl. Fig 6B) and a trend to a lower insulin AUC (Suppl. Fig 6D) during GSIS was observed. The islet Cd bioaccumulation was fond to be markedly lower than in males exposed to the 1 mM CdCl_2_ concentration. We therefore employed a higher Cd exposure concentration of 1mM for future experiments in females.

**Fig 2:**
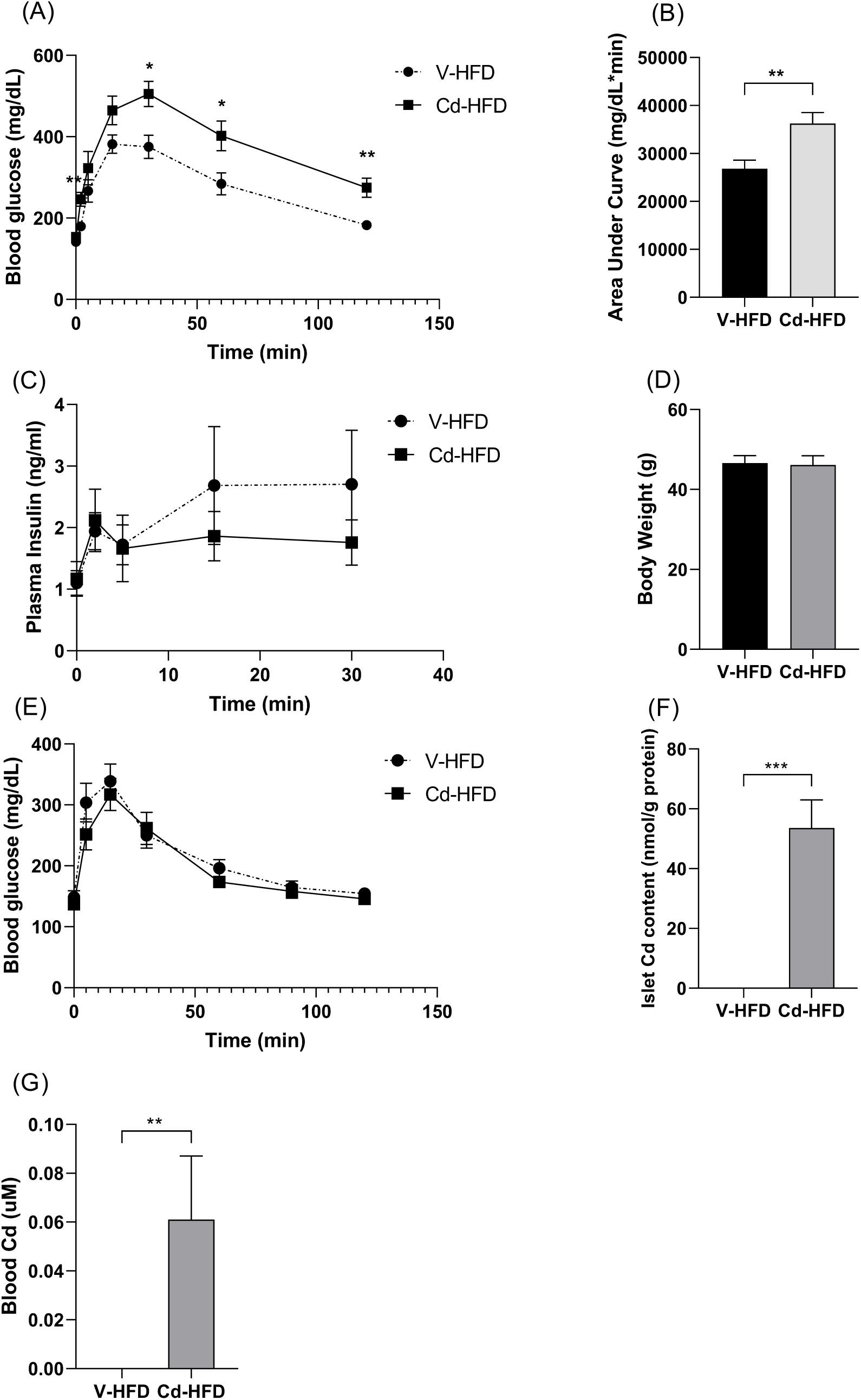
(A) Blood glucose, (B) area under the curve, (C) plasma insulin and (D) fasting body weight during GSIS about 11.4 ± 0.1 weeks into washout phase with HFD in male mice exposed to either V (n=7) or 1mM CdCl_2_ (n=6) for 19.1 ± 0.7 weeks with HFD. (E) Blood glucose during glucose tolerance test 24 weeks into washout phase with HFD in V (n=7) or Cd-exposed (n=6) male mice on HFD (Table 1, Cohort #2). (F) Islet Cd content and (G) blood Cd in male mice exposed to either V (n=6) or 1mM CdCl_2_ (n=6) for 19 weeks with HFD followed by 31.3 ± 0.1 weeks of washout with HFD (Table 1, Cohort #3). V-HFD, Vehicle-exposed followed by washout and HFD; Cd-HFD, Cd-exposed followed by washout and HFD. Data are expressed as mean ± SEM. *p<0.05, **p<0.01, ***p<0.001.

### Islet cadmium bioaccumulation reduces plasma insulin during hyperglycemic clamp without significant changes in *ex vivo* islet insulin secretion

Next, we investigated the effects of islet Cd bioaccumulation on insulin and glucose homeostasis in the setting of insulin resistance. To do this, we exposed age-matched, male mice to either vehicle (n=6) or 0.5mM CdCl_2_ (n=7) in drinking water and fed regular chow for 17 weeks (Table 1, Cohort #5). At the end of 17 weeks Cd exposure, Cd-exposed mice were transitioned to regular vehicle water (i.e. washout phase) with HFD for 12 weeks prior to standard hyperglycemic clamp studies (Ayala, Samuel et al. 2010). In the setting of hyperglycemia, maintained at 250-300mg/dL arterial glucose (Fig. 3A) by continuous glucose infusion (Fig. 3B), Cd-exposed mice on HFD (Cd-HFD) showed lower plasma insulin levels compared to V mice on HFD (V-HFD) at 120 min (p<0.05, Fig. 3C). The decrease in plasma insulin observed in Cd-HFD mice during hyperglycemic clamp occurred in the absence of changes in either plasma c-peptide (Fig. 3D) or body weight (Fig. 3E). To assess Cd accumulation on *ex vivo* islet insulin secretion, we performed islet perifusion and stimulation with glucose and several insulin secretagogues in islets from V-HFD (n=5) and Cd-HFD (n=5) mice. Islet insulin release (Fig. 3F) and total islet insulin content (Fig. 3G) were similar between Cd-HFD and V-HFD control islets in response to 16.7mM glucose, 100uM BMX, 20mM Arginine, 300uM Tolbutamine and 20mM KCl. This suggests that the mechanisms of Cd-induced decrease in plasma insulin observed during hyperglycemic clamp is independent of changes in the rate of insulin secreted per islet or that the *in vivo* milieu is relevant for the Cd’s effect.

**Fig. 3:**
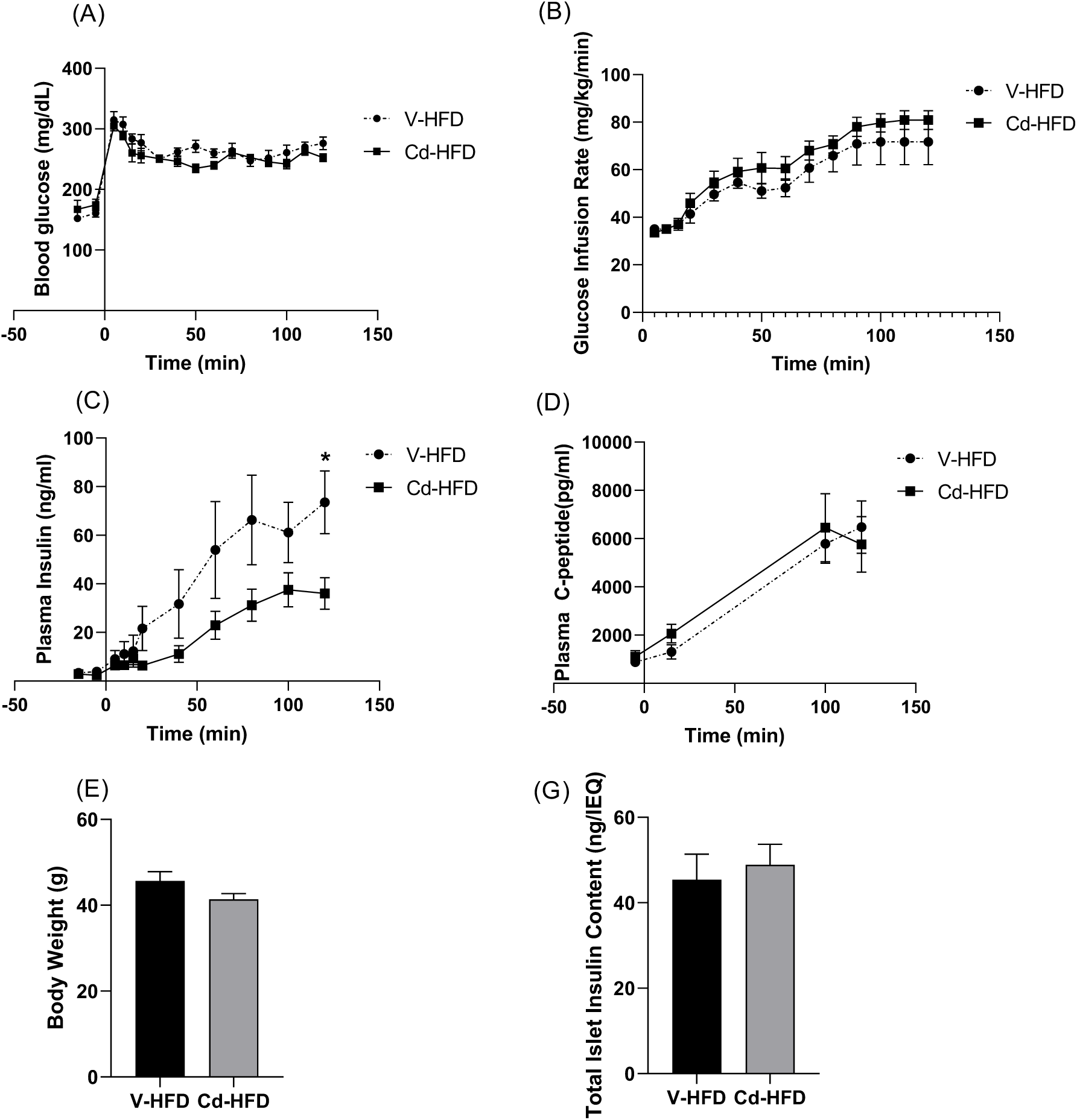
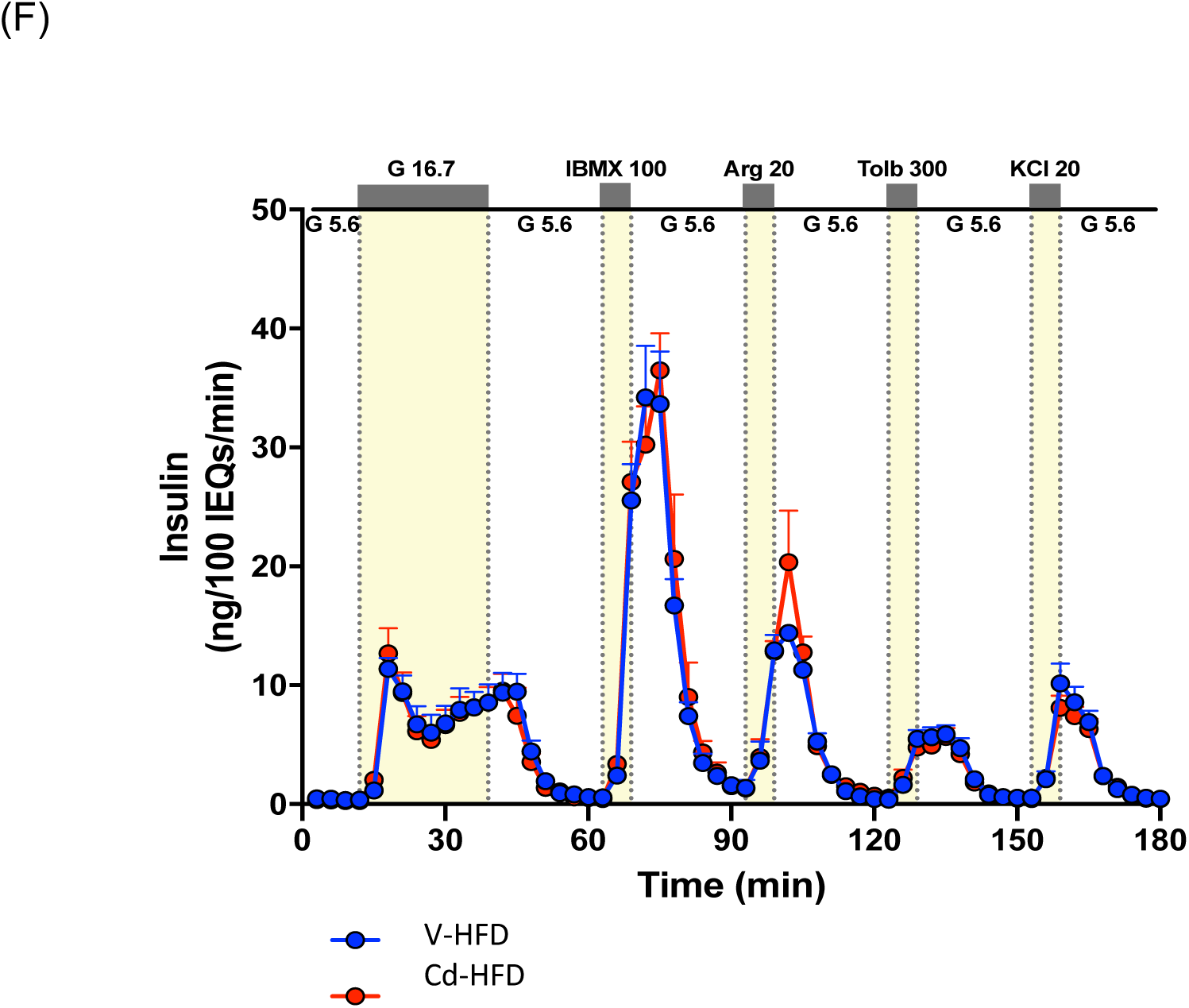
(A) Arterial blood glucose, (B) glucose infusion rate, (C) plasma insulin, (D) plasma c-peptide and (E) body weight during hyperglycemic clamp in mice exposed to either V (n=6) or 0.5mM CdCl_2_ (n=7) for 17 weeks followed by 12 weeks of washout with HFD on V water (Table 1, Cohort #6). (F) Islet insulin secretion in response to high glucose and several insulin secretagogues from V and Cd-exposed mice following 12 weeks of washout with HFD (n=5/group). (G) Total islet insulin content following islet perifusion in V and Cd-exposed mice islets following 12 weeks of washout with HFD (n=5/group). V-HFD, Vehicle-exposed followed by washout and HFD; Cd-HFD, Cd-exposed followed by washout and HFD. Data are expressed as mean ± SEM. *p<0.05.

### Gender dichotomy of cadmium-mediated glycemic effects *in vivo*

Next, we investigated the gender dichotomy of Cd-mediated glycemic effects. To do this, we exposed age-matched, male (Table 1, Cohort 7a) and female (Table 1, Cohort 7b) mice in parallel to either V (n=9-11) or 1mM CdCl_2_ (n=9-10) dose in drinking water for 17.8 ± 0.1 weeks. Cadmium-exposed mice were transitioned to vehicle drinking water following 17.8 ± 0.1 weeks of Cd exposure during the washout phase. Both groups of V and Cd-exposed mice received regular chow during the Cd exposure phase and HFD (42% from fat) during the washout phase. Following the Cd exposure phase and prior to the washout phase, V and Cd-exposed, male mice did not display any differences in whole body glucose clearance (Fig. 4A) and GSIS (Fig. 4B). In contrast, Cd-exposed female mice displayed significantly higher plasma insulin levels 30 minutes after a glucose load compared to their V controls (p<0.05, Fig. 4B). This increase in plasma insulin was not associated with changes in either whole-body glucose clearance (Fig. 4A) or fasting body weight (Fig. 4C). Following 19.1 ± 0.1 weeks into the washout phase with HFD, we did not observe significant differences in glucose values or plasma insulin during GSIS between V-HFD and Cd-HFD males (Figs. 4D-F), a different result compared to prior Cohorts #2 and #3 which received HFD during the Cd exposure phase. In contrast, following the same washout period (19.0 ± 0.8 weeks), we observe a significantly higher whole-body glucose clearance (i.e. lower glucose levels) in Cd-HFD females compared to their V-HFD control counterparts (p<0.05, Fig. 4D and 4E). This increase in whole body glucose clearance in Cd-HFD females was not associated with changes in either plasma insulin (Fig. 4F) or fasting body weight (Fig. 4G). There were also no significant differences in GSIS between Cd-HFD and V-HFD females at 34 weeks into the washout phase (Suppl. Fig 5A-E). Islets from Cd-HFD female mice showed significant islet Cd bioaccumulation (119.3 ± 14.0nmol/g protein) compared to their V-HFD female controls (29.3 ± 17.3nmol/g protein) (p<0.01, Fig. 4H). While islet Cd in V-HFD female controls was significantly lower, it was still markedly higher (29.3 ± 17.3nmol/g protein, p<0.01, Fig. 4H) than V-HFD in males from Cohorts #1, 2, 3 and 5 (Table 1). It is of note that serum creatinine levels in Cd-HFD animals were not significantly different to that of the V-HFD group (Suppl. Table 1) suggesting that this regiment of Cd exposure (1mM CdCl_2_ for 18 weeks followed by washout and HFD for 26 weeks) did not affect kidney function, which is an important consideration when interpreting dysglycemia data given that renal failure results in prolonged half-life of insulin. There were no differences in plasma proinsulin or proinsulin:insulin ratio in selected experiments (data not shown).

**Fig. 4:**
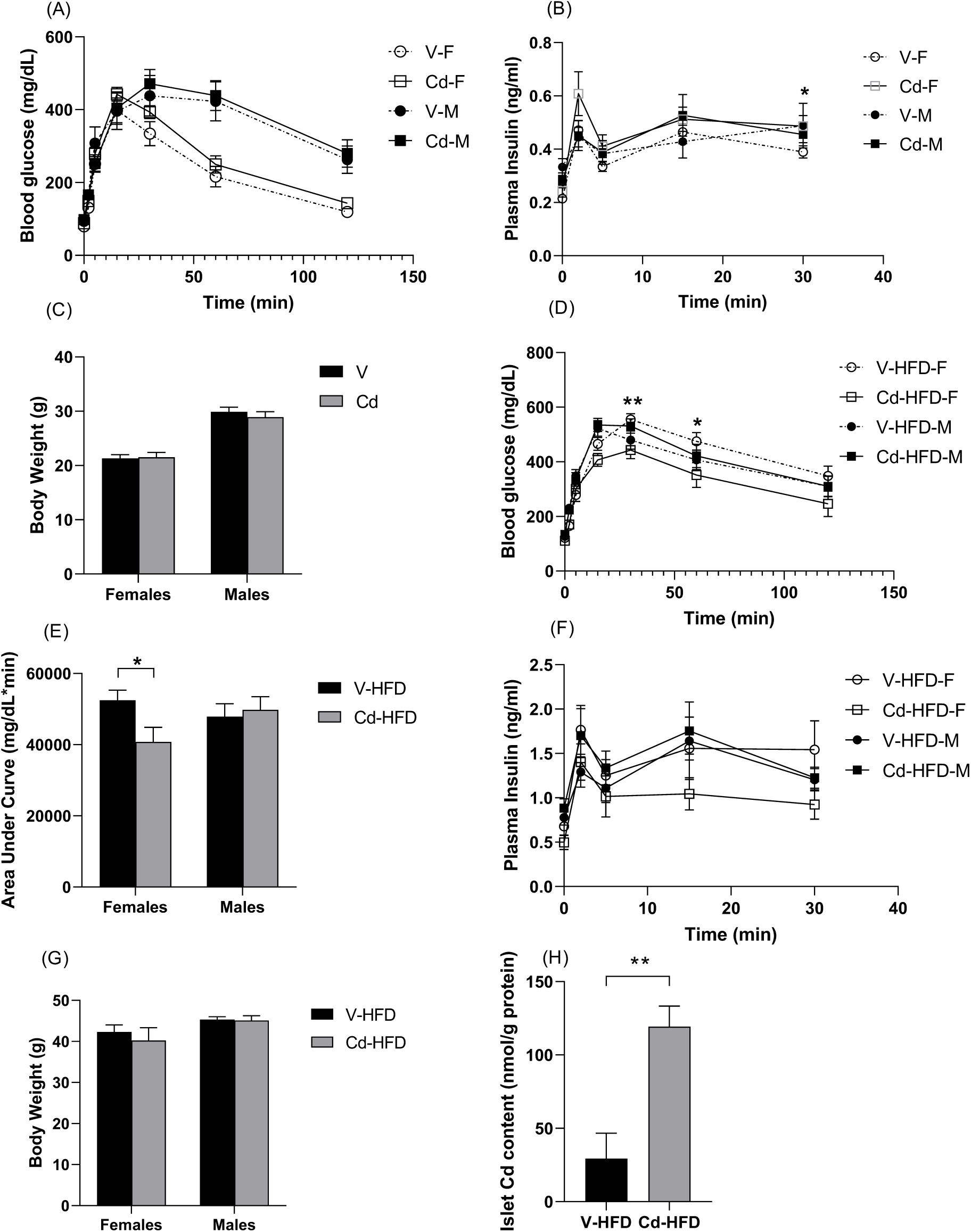
(A) Blood glucose, (B) plasma insulin and (C) fasting body weight during *in vivo* GSIS in age-matched male and female mice exposed to either V (n=10 (males), n=11 (females)) or 1mM CdCl_2_ (n=10/gender) for 17.8 ± 0.1 weeks (Table 1, Cohort #7a-b). (D) Glucose curve and corresponding (E) area under the curve, (F) plasma insulin and (G) fasting body weight during GSIS in age-matched V-HFD and Cd-HFD male (n=9-10/group) and female (n=9-10/group) mice 19.1 ± 0.1 weeks into the washout phase with HFD. (H) Islet Cd content (normalized for total islet protein) at sacrifice in V-HFD (n=9) and Cd-HFD (n=8) female mice. V-F, Vehicle females; Cd-F, Cd-exposed females; V-M, Vehicle males; Cd-M, Cd-exposed males; V-HFD, Vehicle-exposed followed by washout and HFD; Cd-HFD, Cd-exposed followed by washout and HFD. Data are expressed as mean ± SEM. *p<0.05, **p<0.01.

### Linear regression analysis and correlation between islet Cd and plasma insulin

To further evaluate the relationship between Islet Cd bioaccumulation and Cd-mediated dysglycemia in each gender, we performed linear regression analysis between islet Cd content and insulin AUC during *in vivo* GSIS in both males and females. The aim of this analysis is to explore the influence of variations in islet Cd accumulation on plasma insulin given the variation in insulin values seen across cohorts. In cohorts where repeated GSIS was performed, GSIS data closest to the time of islet isolation was chosen. When male and female datasets were pooled and analyzed, we observed a significant gender effect on insulin AUC (p<0.05, Table 2). We therefore repeated the linear regression analysis in males and females separately. We observed that islet Cd content (independent variable) was negatively associated with insulin AUC (dependent variable) during *in vivo* GSIS in males (p<0.05, Table 2) but not in females. There was no correlation between islet Zn and insulin AUC in either males or females (Table 2). The correlation between insulin AUC and islet Cd as well as the gender dichotomy did not change when AUC glucose was included in the model (Suppl. Table 2A). This correlation also did not change in simpler models that included fewer independent variables or islet Cd alone in females (Suppl. Table 2D) but not in males (Suppl. Table 2C). In males, a significant correlation was observed in a model that did not include age between insulin AUC and islet Cd content (Suppl. Table 2C).

**Table 2:** Linear regression analysis between insulin AUC (dependent variable), islet Cd content, islet Zn content and age (independent variable) in males alone (n=33), females alone (n=29) and combined male and female mice (n=62). *p<0.05, ***p<0.001.

|  |  | Model 1 |  |  |  | Model 2 |  |  |  |
| --- | --- | --- | --- | --- | --- | --- | --- | --- | --- |
| Gender | Predictor | Parameter<br>Estimate | LCL<br>Mean | UCL<br>Mean | Pr > \|t\| | Parameter<br>Estimate | LCL<br>Mean | UCL<br>Mean | Pr > \|t\| |
| Males and<br>females | Gender | 5.65 | -8.48 | 19.79 | 0.435 | -17.36 | -33.43 | -1.29 | 0.039* |
|  | Islet Cd | -0.15 | -0.28 | -0.02 | 0.028* | -0.09 | -0.2 | 0.03 | 0.138 |
|  | Islet Zn | 0.0005 | -12X10<sup>-5</sup> | 0.001 | 0.115 | 0.0004 | -21X10<sup>-5</sup> | 0.0009 | 0.217 |
|  | Age | - | - | - | - | 1.36 | 0.75 | 1.98 | <0.001*** |
| Males | Islet Cd | -0.27 | -0.48 | -0.06 | 0.018* | -0.19 | -0.37 | -0.02 | 0.042* |
|  | Islet Zn | 0.0009 | -5 X 10<sup>-6</sup> | 0.002 | 0.061 | 0.0007 | 18X10<sup>-7</sup> | 0.001 | 0.059 |
|  | Age | - | - | - | - | 1.38 | 0.70 | 2.07 | <0.001*** |
| Females | Islet Cd | -0.04 | -0.16 | 0.09 | 0.588 | -0.02 | -0.16 | 0.13 | 0.844 |
|  | Islet Zn | -3X10<sup>-4</sup> | -0.001 | 0.0006 | 0.538 | -44X10<sup>-5</sup> | -0.002 | 0.0006 | 0.435 |
|  | Age | - | - | - | - | 0.68 | -1.85 | 3.2 | 0.604 |

LCL: lower control limits, UCL: upper control limits, Pr > |t| : t-distribution two-tailed p-value.

### *Ex vivo* cadmium accumulation in intact murine and human islets did not result in measurable changes in insulin secretion

To compare the rate of Cd accumulation and functional impact between human and murine islets, we exposed human islets (n=3/group) and mouse islets (n=6/group) to either V or 0.1μM CdCl_2_ for 48 hours in culture and measured islet Cd content. Islets originating from the same donor or mouse were divided equally between parallel exposure conditions. Cadmium-exposed human islets accumulated significantly higher intracellular Cd levels (121.7 ± 25.8nmol/g protein) compared to vehicle control islets (21.1 ± 7.0nmol/g protein) following 48 hours of Cd exposure (p<0.05, Fig. 5A).

**Fig. 5:**
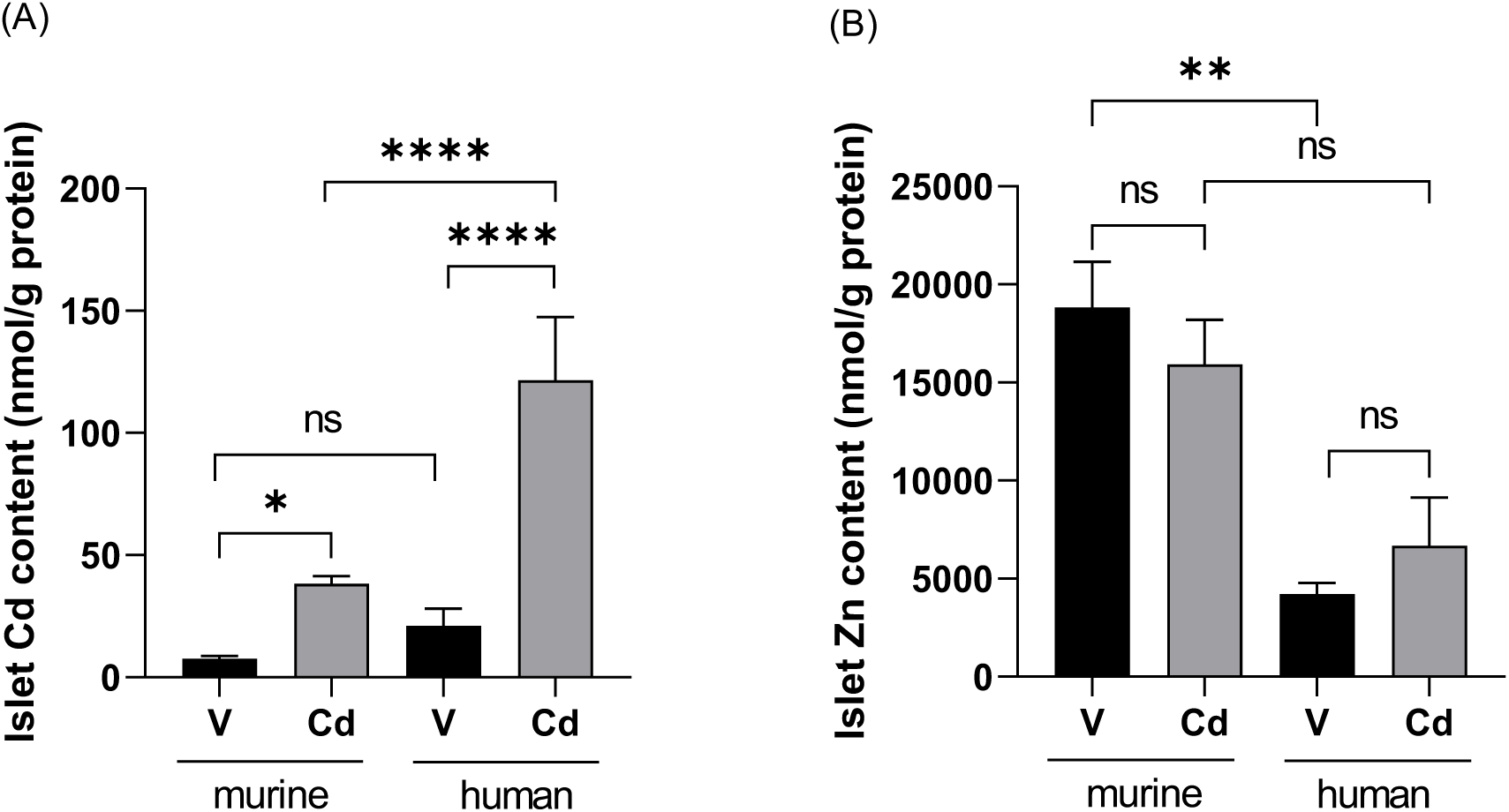
(A) Islet Cd and (B) islet Zn (normalized to total islet protein) in murine (n=6/group) and human islets (n=3/group) exposed to either V or 0.1μM CdCl_2_ for 48 hours. Data are expressed as mean ± SEM. Adjusted p-values from one way ANOVA with post hoc Tukey’s test for multiple comparison. * p<0.05, **p<0.01, ****p<0.0001, ns: not significant.

Cadmium-exposed murine islets also showed significantly higher Cd levels (38.3 ± 3.0nmol/g protein) compared to vehicle control islets (7.6 ± 1.2nmol/g protein) following 48 hours of Cd exposure (p<0.001, Fig. 5A). Cadmium levels in human islets exposed to Cd *ex vivo* was significantly higher compared to mouse islets exposed under the same conditions (p<0.0001, Fig 5A). There were also no significant differences in total islet proinsulin to insulin ratio in response to Cd exposure in mouse islets *ex vivo* (data not shown).

### *Ex vivo* cadmium accumulation did not affect islet Zn content or β-cell Zn turnover

Given that Cd has been shown to interact with many Zn transport and binding proteins (Himeno, Yanagiya et al. 2009, Friedman 2014), we compared islet Zn levels following *ex vivo* (refer to paragraph above) or *in vivo* Cd accumulation in islets. Human islets contained lower Zn levels compared to mouse islets at baseline (p<0.01, Fig 5B), suggesting significant differences in divalent metal handling between human and mouse islets. However, Cd exposure did not significantly affect islet Zn content in either human or mouse islets (Fig 5B). Next, we examined the turnover of Zn following Cd accumulation in immortalized murine β-cell line MIN6. Cell culture, Cd exposure and cell harvest were performed as previously described (El Muayed, Raja et al. 2012). For this, we exposed MIN 6 cells to either V or 0.5μM CdCl_2_ in media for 48 hours. Following 48 hours of CdCl_2_ incubation, media was changed and 10μM ^67^Zn tracer was added to fresh culture media (equivalent to 66% of total Zn in media) and the rate of cellular ^67^Zn uptake was measured at 1, 2 and 4 hours following the start of ^67^Zn tracer exposure. The rate of cellular ^67^Zn uptake was used as a proxy measurement for cellular Zn turnover. We observed that the rate of ^67^Zn accumulation were similar between V and Cd-exposed MIN6 cells, suggesting that cellular Cd accumulation *in vitro* did not affect cellular Zn turnover (Suppl. Fig. 2). Overall, islet Zn content was not significantly different between Cd-HFD mice and V-HFD controls from any of our animal cohorts (Table 1).

## Discussion

Here, we report the development of a chronic, oral Cd exposure mouse model that results in islet Cd bioaccumulation that is comparable to that found in human islets from the general US population (Wong, Allen et al. 2017). In males, exposure to 1mM CdCl_2_ in drinking water for 19 weeks followed by washout with HFD resulted in islet Cd bioaccumulation that is associated with decrease glucose tolerance at 11 weeks and decrease plasma insulin at 28 weeks into the washout phase in the absence of changes in *ex vivo* islet insulin secretion, body weight or renal function. In contrast, Cd exposure in female mice to the same conditions was associated with islet Cd bioaccumulation and elevated plasma insulin during GSIS without changes in whole body glucose clearance immediately following Cd exposure (prior to washout phase). However, following the washout phase with HFD, Cd-HFD females displayed lower systemic glucose clearance during GSIS without changes in either plasma insulin or body weight despite achieving comparable islet Cd concentrations. A relatively wide variation in islet Cd accumulation was observed between individual mouse. This also included a relatively high level of Cd in female vehicle-exposed islets in cohort 7b. It is likely that variation in consumption of CdCl_2_-containing water as well as varied oral bioavailability and tissue distribution are factors contributing to this. Additionally, variations in the content of Cd and other metals in rodent chows may have contributed to the observed variations.

Results from the regression analysis showed the close relationship between islet Cd plasma insulin levels during *in vivo* glucose-stimulated insulin secretion experiments, providing additional evidence for the functional impact of Cd accumulation at environmentally relevant concentrations on *in vivo* beta-cell function. At the same time, this relationship indicates that variations in the observed effect of Cd exposure across cohorts and within cohorts are at least partially explained by variations in islet Cd accumulation. There are several benefits to our oral exposure-washout *in vivo* mouse model. Firstly, it ameliorated the anorexigenic effects of Cd, thereby resulting in a similar mean body weight gain between V and Cd-exposed animals throughout the study. The *in vivo* anorexigenic effects of Cd was mitigated with the washout phase following Cd exposure in our model. Secondly, this oral exposure-washout regiment achieves target tissue (pancreatic islets) Cd bioaccumulation at concentrations comparable to those found in human islets while reducing systemic blood Cd concentrations at the end of the washout phase and minimizing generalized systemic toxic effects. Islet Cd range in Cd-exposed mice is consistent with islet Cd range that we have previously reported in human islets (4.4 - 94.7nmol/g protein) (Wong, Allen et al. 2017), hence making this *in vivo* experimental Cd dose relevant in the context of human environmental Cd exposure. Furthermore, the continued oral Cd exposure through drinking water was employed to resemble environmental exposure routes seen in the general population. The oral Cd dose of 1mM was chosen to compensate for low oral Cd bioavailability and life-long tissue accumulation as reported in humans (Jarup, Rogenfelt et al. 1983, Schilderman, Moonen et al. 1997).

Several epidemiological studies have described contradictory findings between Cd exposure and dysglycemia. Some have reported a statistically significant, positive association between Cd exposure and dysglycemia (Schwartz, II’yasova et al. 2003, Wallia, Allen et al. 2014, Nie, Wang et al. 2016) while others have reported the lack of a statistically significant association (Barregard, Bergstrom et al. 2013, Moon 2013, Menke, Guallar et al. 2016). Discrepancies in observations could be attributed to differences in study design, study power, sample analyzed (blood or urine), genetics and population exposure levels. Inconsistencies between Cd exposure and dysglycemia have also been reported in Cd-induced diabetic animal models. Some studies have shown that *in vivo* Cd exposure resulted in hyperglycemia and β-cell dysfunction (Chang, Hsu et al. 2013, Trevino, Waalkes et al. 2015, Huang, Kuo et al. 2019, Fitzgerald, Olsen et al. 2020, Hong, Xu et al. 2021) while others have demonstrated normal glucose homeostasis with modest decrease in insulin secretion following Cd exposure *in vivo* (Li, Li et al. 2019). Trevino *et al* (2015) and Fitzgerald *et al* (2020) used a lower Cd dose of 32.5ppm and 0.6ppm in their exposure rat models respectively. Animal strain, dose, duration and route of administration of Cd should be considered when interpreting data reported in Cd-induced animal models of dysglycemia. Additionally, short duration Cd exposure may result in acute systemic toxic effects especially with elevated blood Cd levels in the absence of a washout period. Given the diverse intracellular targets of Cd, each with a different concentration-effect relationship, it is not surprising that Cd effects that are observed at a whole cell and organism level are strongly dependent on exposure duration and tissue concentration.

Pancreatic β-cells accumulate Cd in a dose- and time-dependent manner *in vitro* (El-Muayed, Raja et al. 2012). Measurable levels of Cd have also been detected in human islets (Wong, Allen et al. 2017). However, the molecular mechanisms of Cd^2+^ transport into β-cells are largely unknown. DMT1 (Lortz, Schroter et al. 2014), ZnT (Kambe, Narita et al. 2002, Chimienti, Favier et al. 2005, Cai, Kirschke et al. 2018) and ZIP (Lawson, Maret et al. 2017) are metal transport proteins that are abundantly expressed in pancreatic β-cells. A known iron transporter, DMT1 has been reported to also effectively transport several other divalent metals such as Fe^2+^, Mn^2+^, Ni^2+^, Co^2+^and Cd^2+^ (Tallkvist, Bowlus et al. 2001, Arredondo, Munoz et al. 2003, Garrick, Dolan et al. 2003). ZIP and ZnT are known to facilitate Zn^2+^ entry into and exit from the cytosol respectively (Lemaire, Chimienti et al. 2012, Lawson, Maret et al. 2017). In addition to transporting the divalent cation Zn^2+^, some ZIP class transporters can transport Cu^2+^, Ni^2+^ (Antala and Dempski 2012) and Mn^2+^ (Winslow, Limesand et al. 2020). The capacity to transport Cd effectively has been shown for ZIP8, ZIP14 and ZnT1 (Dalton, He et al. 2005, Ohana, Sekler et al. 2006, Girijashanker, He et al. 2008). DMT-1, ZIP14 and ZIP8 are highly expressed in murine and human islets (Mohanasundaram, Drogemuller et al. 2011, Hansen, Tonnesen et al. 2012, Coffey and Knutson 2017, Lawlor, George et al. 2017, Wong, Wang et al. 2021).

We have previously shown that the accumulation of Cd in β-cells at environmentally relevant levels leads to impaired β-cell function *ex vivo* (El-Muayed, Raja et al. 2012). These experiments were carried out in dispersed murine islets as well as in an immortalized murine β-cell line, MIN6. In contrast, our current experiments did not show any differences in *ex vivo* function of intact islets following either *in vivo* or *ex vivo* Cd accumulation. It is likely that this difference is due to a protective effect of an intact islet structure. We also observed a higher rate of *ex vivo* Cd accumulation in human islets compared to mouse islets, possibly due to differences in the composition and function of metal transporters (especially divalent metal transporters) between human and mouse islets. This observation highlights the value of assessing tissue level Cd concentrations when evaluating its toxic effects. Accumulation of Cd *in vivo* have been associated with hyperglycemia and decrease glucose-stimulated insulin secretion in a model of shorter term exposure (Fitzgerald, Olsen et al. 2020). The addition of a washout phase in our current model provides the added advantage of mitigating acute toxic effects of Cd, thereby facilitating a better correlation between target tissue Cd levels and its resulting toxic effects.

The underlying molecular mechanisms of intracellular Cd bioaccumulation on β-cell cytotoxicity and function are not well studied. Chang et al. (2013) showed that intracellular Cd accumulation *in vitro* resulted in β-cell apoptosis that is mediated by the JNK-regulated, mitochondria-dependent pathway (Chang, Hsu et al. 2013). Cadmium-mediated pancreatic β-cell death was also shown to be mediated by Ca^2+^-dependent, JNK activation and CHOP-related signaling pathways (Huang, Kuo et al. 2019). It has also been hypothesized that Cd may interfere with the trafficking of Zn in β-cells given the high Zn turnover present in β-cells and the overlap between Cd and Zn transport pathways. However, the lack of change in islet Zn concentration in response to Cd accumulation provides evidence against this hypothesis, at least at the whole cell level. However, this does not rule out changes in Zn distribution within cells at the protein or organelle level in response to Cd accumulation. Taken together, our current findings and published literature indicate that accumulation of Cd in pancreatic islets at environmentally relevant concentrations that are in the range found in human islets (Wong, Allen et al. 2017) is cytotoxic to β-cells, thus resulting in Cd-mediated β-cell changes that consequently affects total insulin secretory capacity *in vivo*. Therefore, the current study adds to prior epidemiologic and experimental evidence that members of the general population are exposed to environmental Cd levels that are likely to contribute to β-cell failure, thereby increasing the risk for type 2 diabetes. Consequently, the diabetogenic effect of Cd should be taken into account when setting Cd exposure limits.

Cadmium-mediated molecular mechanisms *in vivo* and *ex vivo* are likely distinct from one another which, in turn, could affect Cd toxicology differently under both conditions. This may be due to the uptake, organification and toxicokinetic properties of Cd. *In vivo*, Cd is primarily bound to albumin or metallothionein (Jin, Nordberg et al. 1986, Nordberg 2004). *Ex vivo*, the primary form of Cd in culture is Cd^2+^. Recently, we reported differences in Cd-mediated islet transcriptome changes between *in vivo-* and *ex viv*o-exposed islets where we did not find any overlap in Cd-induced differentially expressed genes in both groups (Wong, Wang et al. 2021). While our model shows several advantages over other approaches, several drawbacks have to be acknowledged. Firstly, these experiments are time-consuming and resource-intensive. Secondly, the observed variation in islet Cd accumulation and resulting variations in observed changes in plasma insulin presents a drawback. Despite this, we believe this model to be a good tool for studying the degree and mechanisms underlying the diabetogenic effect of Cd and other persistent toxicants that accumulate in insulin-producing islets.

Here, we report the development of a chronic, oral Cd exposure mouse model that results in islet Cd bioaccumulation that is relevant in the context of environmental Cd exposure in humans. Our *in vivo* Cd exposure-washout mouse model showed islet Cd bioaccumulation that is associated with subtle and complex Cd-mediated effects on glucose clearance and plasma insulin levels. Further research examining the relationship between Cd exposure, islet Cd bioaccumulation, dysglycemia and their underlying mechanisms are needed to understand the effects of Cd-mediated β-cell dysfunction.

## Acknowledgements

ICP-MS metal analysis was performed at Quantitative Bio-elemental Imaging Center, Northwestern University that is supported by NASA Ames Research Center (NNA04CC36G). The authors thank the Vanderbilt MMPC Core Facility (U24 DK059637) for performing hyperglycemic clamps and islet perifusion studies.

## Authors’ contributions

Winifred Wong: Conceptualization, Investigation, Validation, Formal analysis, Writing – original draft, review and editing. Janice Wang: Investigation, Validation, Writing-review and editing. Matthew Meyers: Investigation. Nathan Wang: Investigation. Rebecca Sponenburg: Investigation. Norrina Allen: Formal analysis. Joshua Edwards: Writing – review and editing. Malek El-Muayed: Conceptualization, Investigation, Formal analysis, Funding acquisition, Writing-review and editing. All authors have read and approved the final manuscript.

### Supplemental Figures

**Suppl. Fig. 1:**
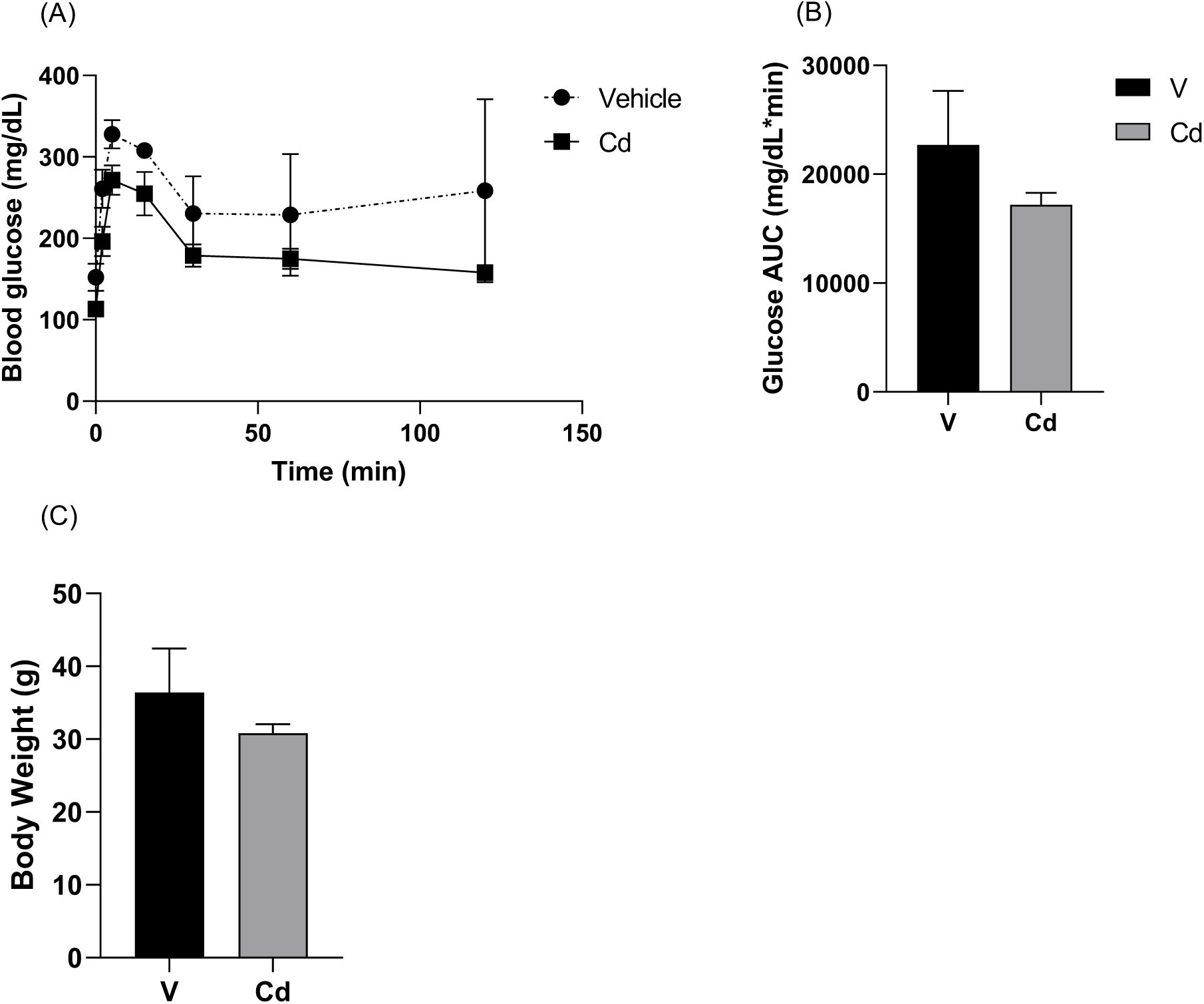
(A) Glucose tolerance test and corresponding (B) area under the curve (AUC) and (C) fasting body weight in mice exposed to either V (n=3) or 1mM CdCl_2_ (n=5) in drinking water for 12 months. Data are expressed as mean ± SEM.

**Suppl. Fig. 2:**
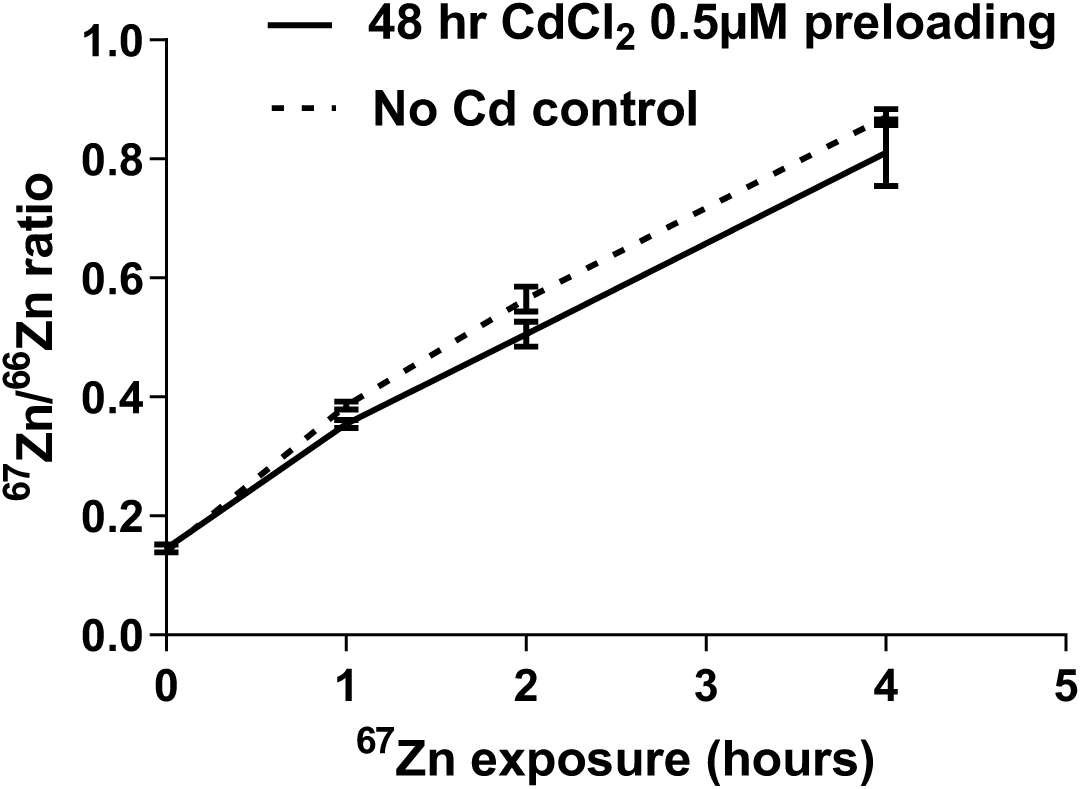
Rate of non-radioactive, rare Zn isotope, ^67^Zn uptake as a proxy for Zn turnover in MIN6 cells following exposure to either V or 0.5µM CdCl_2_ for 48 hours (n=2). Data are expressed as mean ± SEM.

**Suppl. Fig 3:**
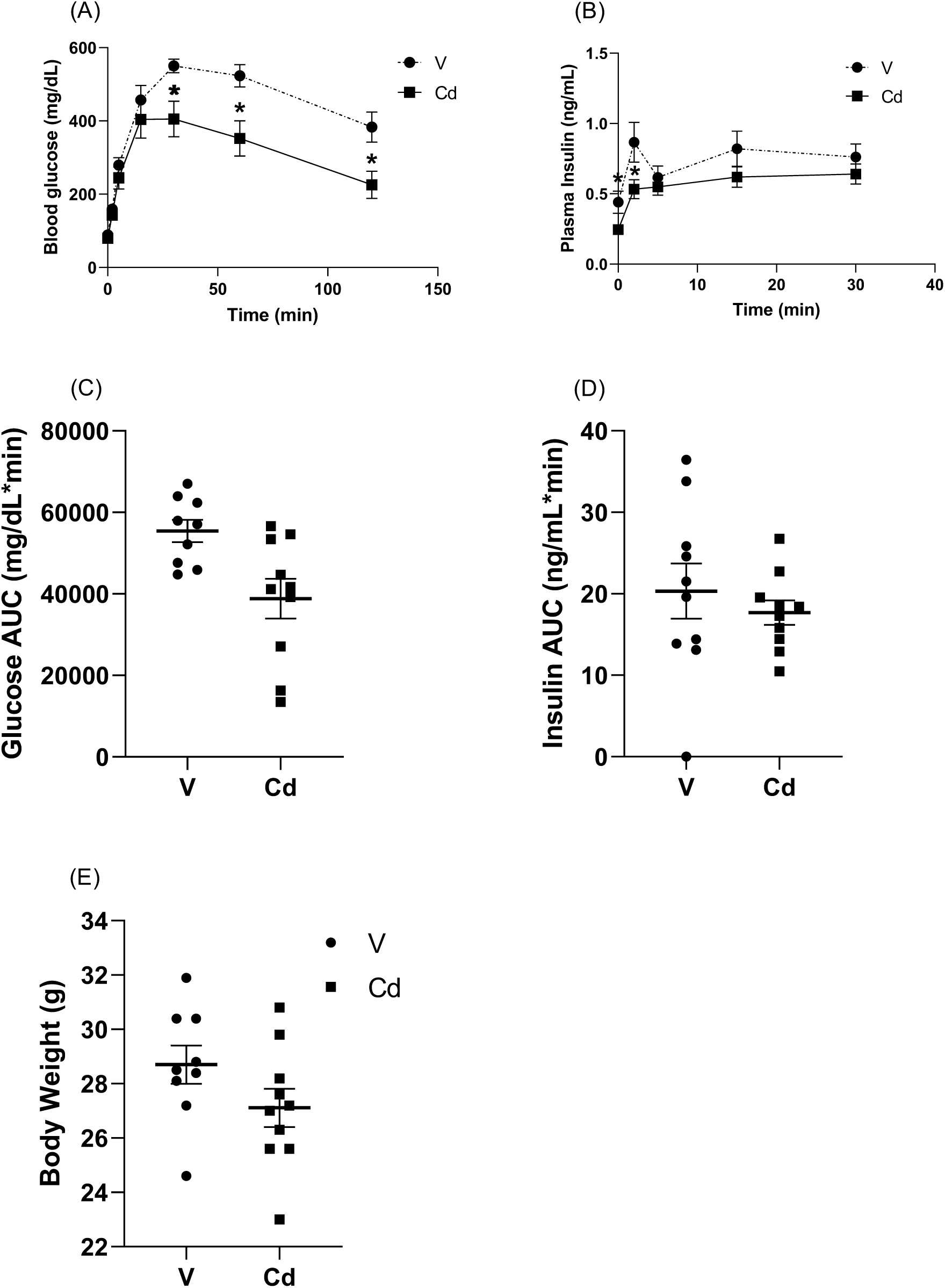
(A) Blood glucose, (B) plasma insulin, (C) AUC glucose, (D) AUC insulin and (E) fasting body weight during *in vivo* GSIS in male mice exposed to either V (n=9) or 1mM CdCl_2_ (n=10) for 19 weeks (Table 1, Cohort #4). GSIS was performed at the end of the Cd exposure phase and prior to the commencement of the washout phase. Data are expressed as mean ± SEM. *p<0.05.

**Suppl. Fig 4:**
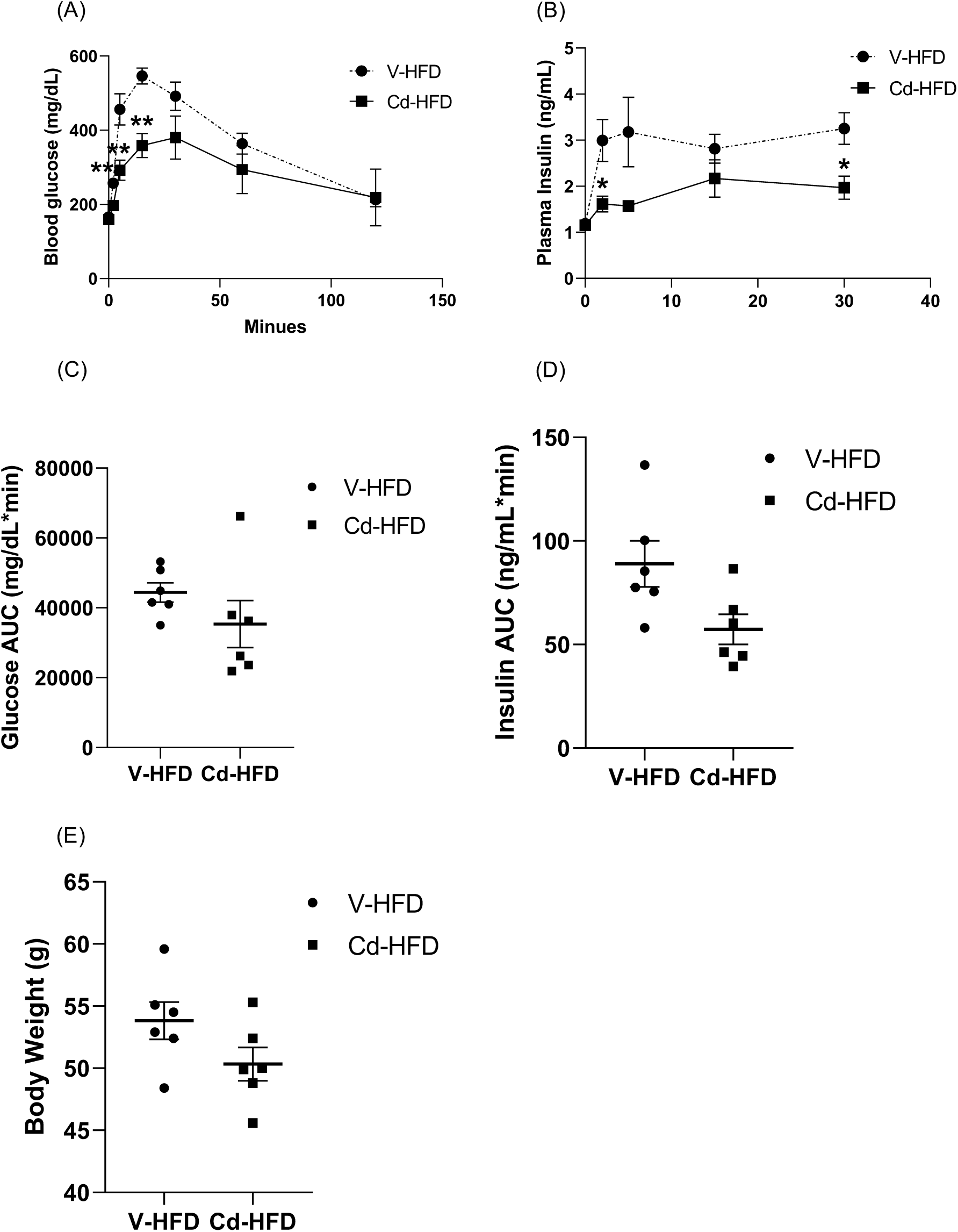
(A) Blood glucose, (B) plasma insulin, (C) AUC glucose, (D) AUC insulin and (E) fasting body weight during *in vivo* GSIS in male mice exposed to either V (n=6) or 1mM CdCl_2_ (n=6) for 19 weeks followed by 28.9 weeks washout phase (table 1, Coort #3). Mice were given HFD at both the Cd exposure and washout phases. V-HFD, V-exposed group given HFD; Cd-HFD, Cd-exposed group given HFD. Data are expressed as mean ± SEM. **p<0.01.

**Suppl. Fig 5:**
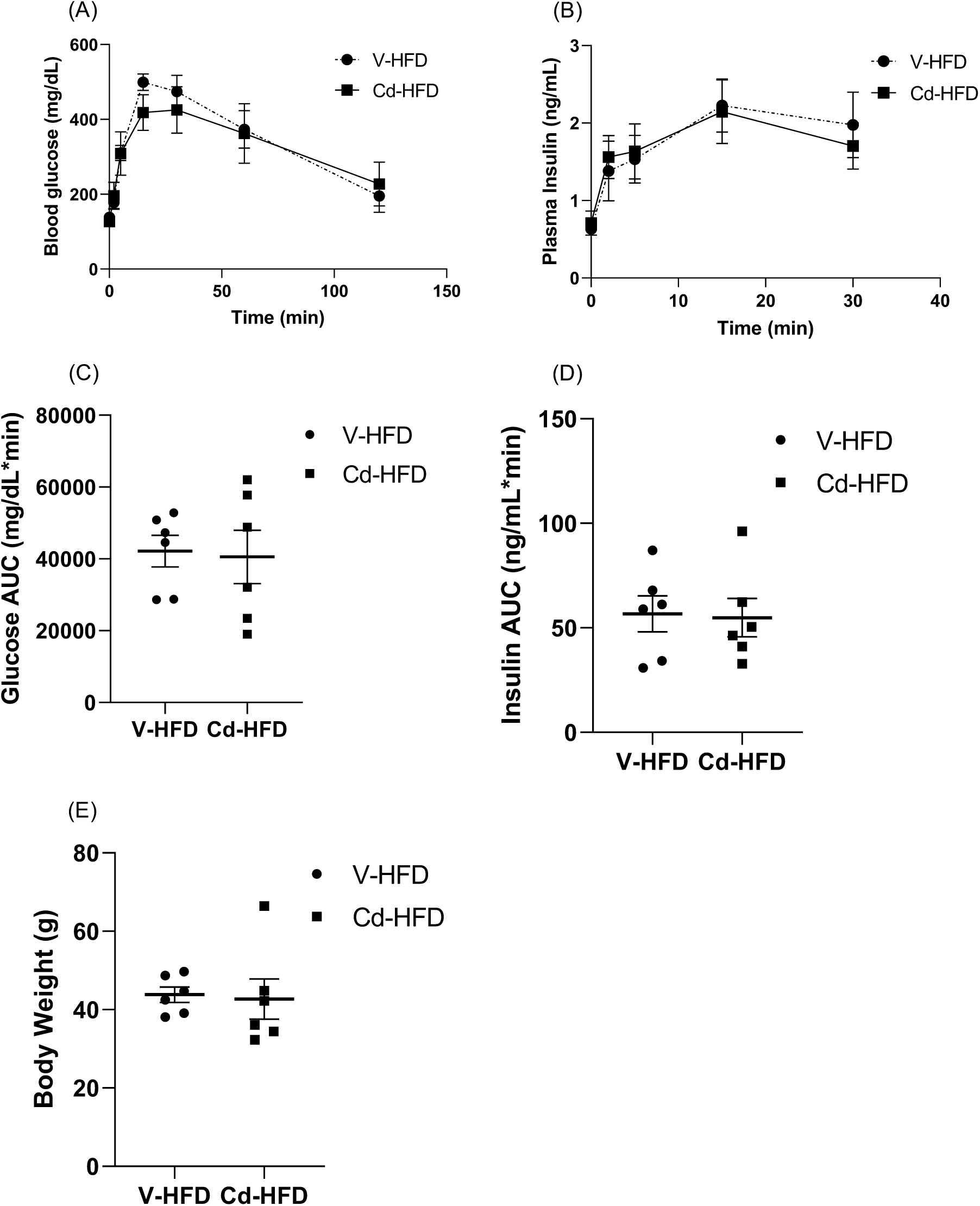
(A) Blood glucose, (B) plasma insulin, (C) AUC glucose, (D) AUC insulin and (E) fasting body weight during *in vivo* GSIS in female mice exposed to either V (n=6) or 1mM CdCl_2_ (n=6) for 18 weeks followed by washout phase for 34 weeks with HFD (Table 1, Cohort #7b). V-HFD, V-exposed followed by washout and HFD; Cd-HFD, Cd-exposed followed by washout and HFD. Data are expressed as mean ± SEM.

**Suppl. Fig 6:**
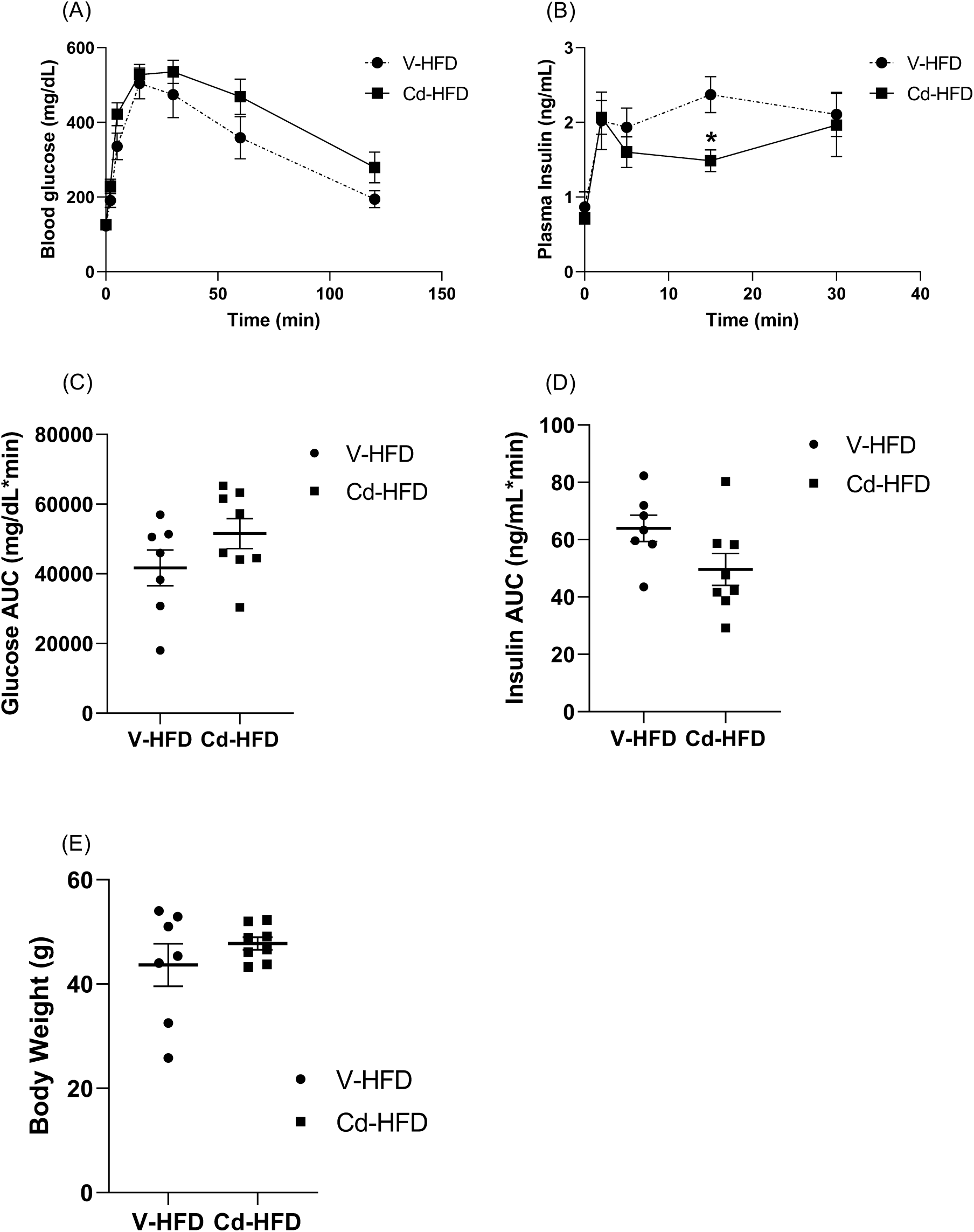
(A) Blood glucose, (B) plasma insulin, (C) AUC glucose, (D) AUC insulin and (E) fasting body weight during *in vivo* GSIS in female mice exposed to either V (n=7) or 1mM CdCl_2_ (n=8) for 18 weeks followed by 36 weeks washout phase (without Cd) (Table 1, Cohort #4). Mice were given HFD at both the Cd exposure and washout phases. V-HFD, V-exposed group given HFD; Cd-HFD, Cd-exposed group given HFD. Data are expressed as mean ± SEM. *p<0.05.

### Supplemental Table

**Suppl. Table 1:**
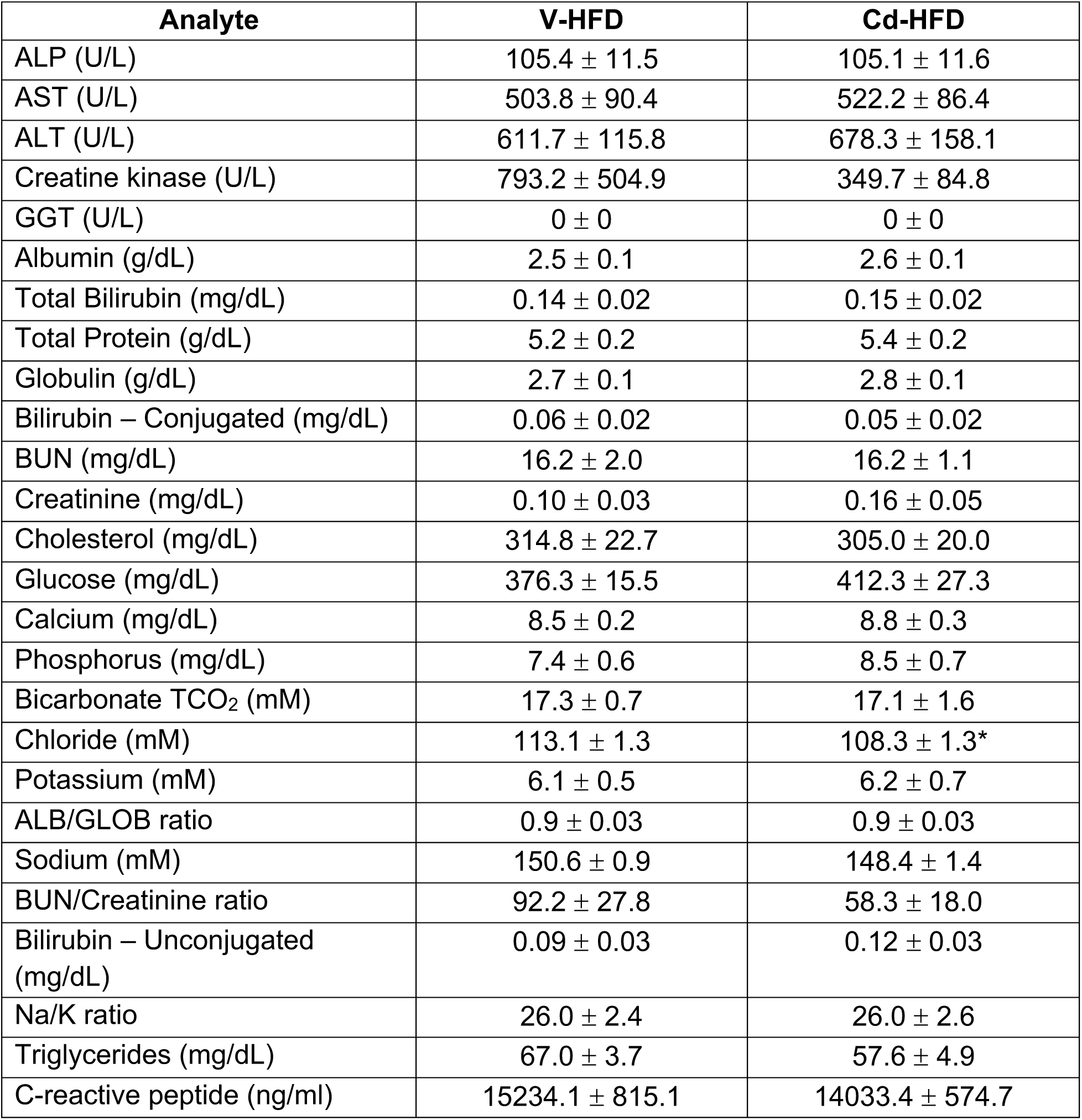
Serum Clinical Chemistry from male mice that were treated with either V (n=9) or 1mM CdCl_2_ (n=10) for 18 weeks followed by 26 weeks HFD thereafter. V-HFD, V-exposed followed by washout with HFD mice; Cd-HFD, Cd-exposed followed by washout with HFD mice. Data are expressed as mean ± SEM. *p<0.05.

**Suppl. Table 2A:**
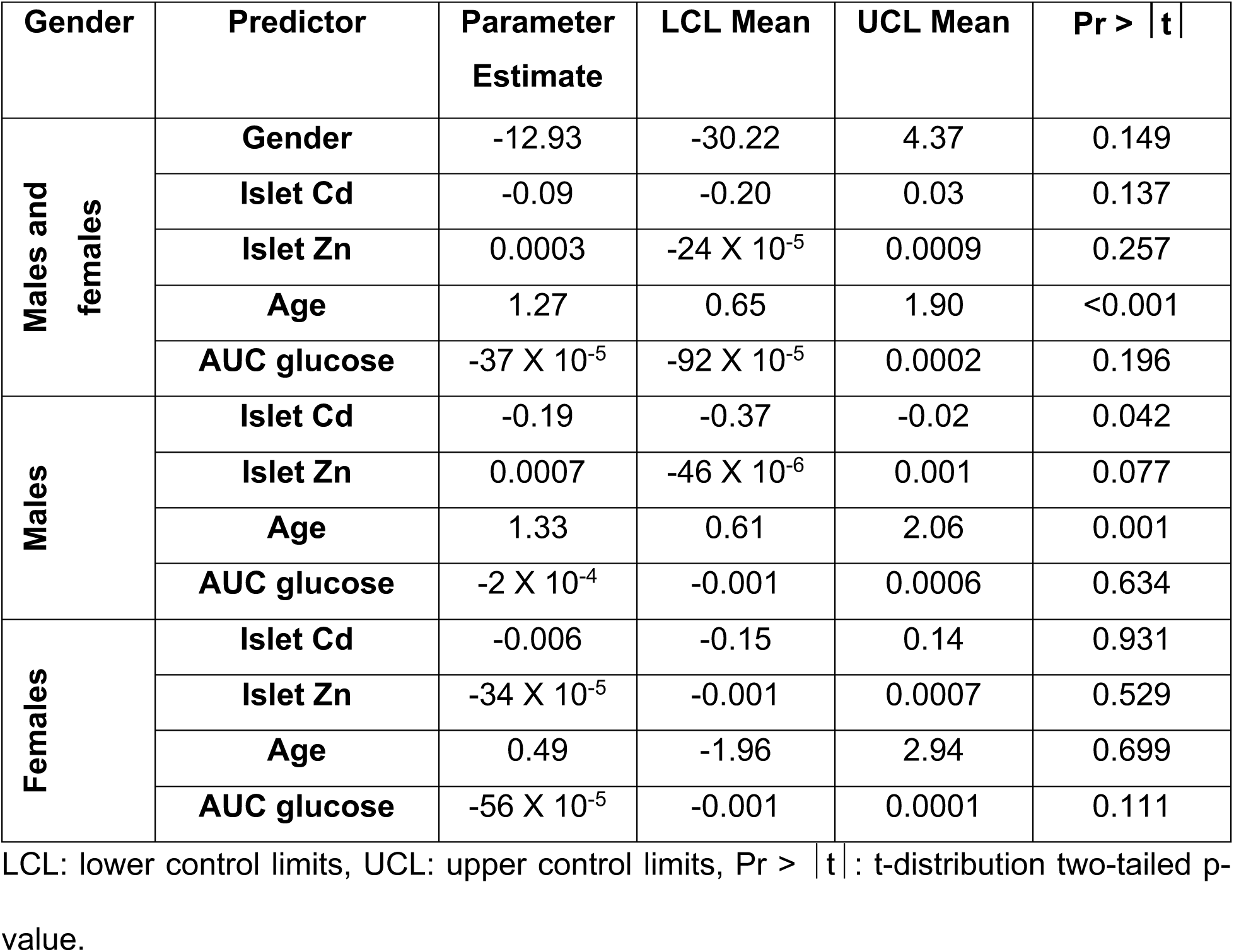
Linear regression analysis between insulin AUC during GSIS (dependent variable) and islet Cd content, islet Zn content, age and glucose AUC as independent variables in males alone (n=33), females alone (n=29) and combined (n=62).

**Suppl. Table 2B:** Linear regression model between insulin AUC during GSIS (dependent variable) and islet Cd content with successive addition of gender, islet Zn content, age and AUC glucose as independent variables in males and females combined (n=62).

|  | Model |  |  |  |  |  |  |  |  |  |  |  |
| --- | --- | --- | --- | --- | --- | --- | --- | --- | --- | --- | --- | --- |
|  | MODEL1 |  |  |  | MODEL2 |  |  |  | MODEL3 |  |  |  |
|  | Parameter Estimate | LCLmean | UCLmean | Pr > \|t\| | Parameter Estimate | LCLmean | UCLmean | Pr > \|t\| | Parameter Estimate | LCLmean | UCLmean | Pr > \|t\| |
| Predictor |  |  |  |  |  |  |  |  |  |  |  |  |
| Islet_Cd | -0.14 | -0.26 | -0.006 | 0.044 | -0.14 | -0.27 | -0.007 | 0.043 | -0.15 | -0.28 | -0.02 | 0.028 |
| female | . | . | . | . | 3.72 | -10.40 | 17.84 | 0.607 | 5.65 | -8.48 | 19.79 | 0.436 |
| Islet_Zn | . | . | . | . | . | . | . | . | 0.0005 | -12E-5 | 0.001 | 0.115 |
| Age | . | . | . | . | . | . | . | . | . | . | . | . |
| AUC_glucose | . | . | . | . | . | . | . | . | . | . | . | . |

|  | Model |  |  |  |  |  |  |  |
| --- | --- | --- | --- | --- | --- | --- | --- | --- |
|  | MODEL4 |  |  |  | MODEL5 |  |  |  |
|  | Parameter Estimate | LCLmean | UCLmean | Pr > \|t\| | Parameter Estimate | LCLmean | UCLmean | Pr > \|t\| |
| Predictor |  |  |  |  |  |  |  |  |
| Islet_Cd | -0.09 | -0.20 | 0.03 | 0.138 | -0.09 | -0.20 | 0.03 | 0.137 |
| female | -17.36 | -33.43 | -1.29 | 0.039 | -12.93 | -30.22 | 4.37 | 0.149 |
| Islet_Zn | 0.0004 | -21E-5 | 0.0009 | 0.217 | 0.0003 | -24E-5 | 0.0009 | 0.257 |
| Age | 1.36 | 0.75 | 1.98 | <.001 | 1.27 | 0.65 | 1.90 | <.001 |
| AUC_glucose | . | . | . | . | -37E-5 | -92E-5 | 0.0002 | 0.196 |

LCL: lower control limits, UCL: upper control limits, $Pr > |t|$ : t-distribution two-tailed p-value.

**Suppl. Table 2C:** Linear regression model between insulin AUC during GSIS (dependent variable) and islet Cd content with successive addition of islet Zn content, age, and AUC glucose as independent variables in males (n=33).

| Predictor | Model |  |  |  |  |  |  |  |  |  |  |  |
| --- | --- | --- | --- | --- | --- | --- | --- | --- | --- | --- | --- | --- |
|  | MODEL1 |  |  |  | MODEL2 |  |  |  | MODEL3 |  |  |  |
|  | Parameter Estimate | LCLmean | UCLmean | Pr > \|t\| | Parameter Estimate | LCLmean | UCLmean | Pr > \|t\| | Parameter Estimate | LCLmean | UCLmean | Pr > \|t\| |
| Islet_Cd | -0.27 | -0.48 | -0.06 | 0.018 | -0.19 | -0.37 | -0.02 | 0.042 | -0.19 | -0.37 | -0.02 | 0.042 |
| Islet_Zn | 0.0009 | -5E-6 | 0.002 | 0.061 | 0.0007 | 18E-7 | 0.001 | 0.059 | 0.0007 | -46E-6 | 0.001 | 0.077 |
| Age | . | . | . | . | 1.38 | 0.70 | 2.07 | <.001 | 1.33 | 0.61 | 2.06 | 0.001 |
| AUC_glucose | . | . | . | . | . | . | . | . | -2E-4 | -0.001 | 0.0006 | 0.634 |

LCL: lower control limits, UCL: upper control limits, $Pr > |t|$ : t-distribution two-tailed p-value.

**Suppl. Table 2D:** Linear regression model between insulin AUC during GSIS (dependent variable) and islet Cd content with successive addition of islet Zn content, age and AUC glucose as independent variables in females (n=29).

| Predictor | Model |  |  |  |  |  |  |  |  |  |  |  |
| --- | --- | --- | --- | --- | --- | --- | --- | --- | --- | --- | --- | --- |
|  | MODEL1 |  |  |  | MODEL2 |  |  |  | MODEL3 |  |  |  |
|  | Parameter Estimate | LCLmean | UCLmean | Pr > \|t\| | Parameter Estimate | LCLmean | UCLmean | Pr > \|t\| | Parameter Estimate | LCLmean | UCLmean | Pr > \|t\| |
| Islet_Cd | -0.04 | -0.16 | 0.09 | 0.588 | -0.02 | -0.16 | 0.13 | 0.844 | -0.006 | -0.15 | 0.14 | 0.931 |
| Islet_Zn | -3E-4 | -0.001 | 0.0006 | 0.538 | -44E-5 | -0.002 | 0.0006 | 0.435 | -34E-5 | -0.001 | 0.0007 | 0.529 |
| Age | . | . | . | . | 0.68 | -1.85 | 3.20 | 0.604 | 0.49 | -1.96 | 2.94 | 0.699 |
| AUC_glucose | . | . | . | . | . | . | . | . | -56E-5 | -0.001 | 0.0001 | 0.111 |

LCL: lower control limits, UCL: upper control limits, $Pr > |t|$ : t-distribution two-tailed p-value.

**Suppl. Table 3:**
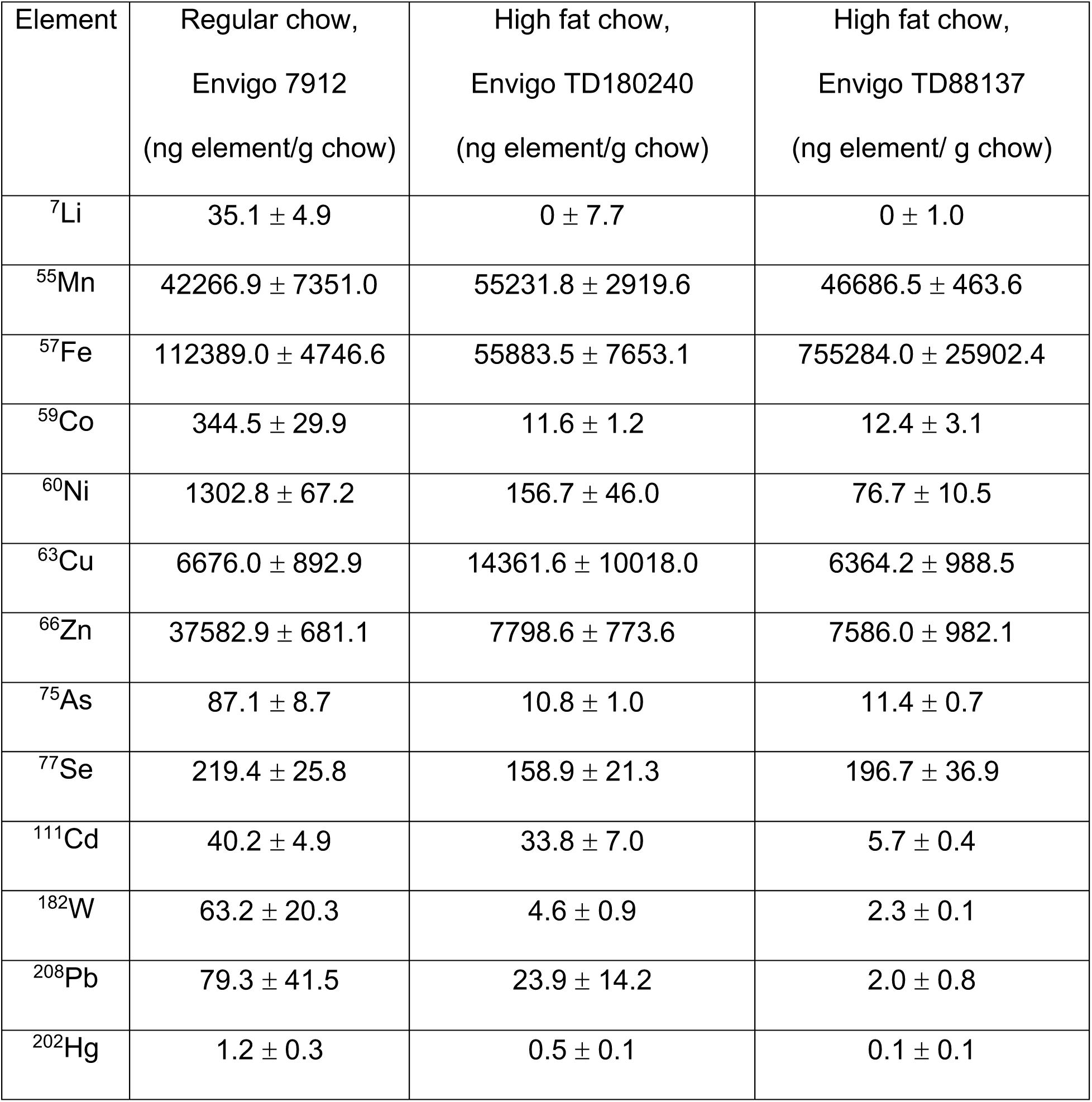
Metal analysis for regular and high fat chow by ICP-MS normalized to chow weight (n=3/group). Data are expressed as mean ± SEM.

## References

1. Afridi, H. I., T. G. Kazi, N. Kazi, M. K. Jamali, M. B. Arain, N. Jalbani, J. A. Baig and R. A. Sarfraz (2008). “Evaluation of status of toxic metals in biological samples of diabetes mellitus patients.” Diabetes Res Clin Pract 80(2): 280–288.

2. Antala, S. and R. E. Dempski (2012). “The human ZIP4 transporter has two distinct binding affinities and mediates transport of multiple transition metals.” Biochemistry 51(5): 963–973.

3. Arredondo, M., P. Munoz, C. V. Mura and M. T. Nunez (2003). “DMT1, a physiologically relevant apical Cu1+ transporter of intestinal cells.” Am J Physiol Cell Physiol 284(6): C1525–1530.

4. Ayala, J. E., V. T. Samuel, G. J. Morton, S. Obici, C. M. Croniger, G. I. Shulman, D. H. Wasserman, O. P. McGuinness and N. I. H. M. M. P. C. Consortium (2010). “Standard operating procedures for describing and performing metabolic tests of glucose homeostasis in mice.” Dis Model Mech 3(9-10): 525–534.

5. Banerjee, M., P. Vats, A. S. Kushwah and N. Srivastava (2019). “Interaction of antioxidant gene variants and susceptibility to type 2 diabetes mellitus.” Br J Biomed Sci 76(4): 166–171.

6. Barregard, L., G. Bergstrom and B. Fagerberg (2013). “Cadmium exposure in relation to insulin production, insulin sensitivity and type 2 diabetes: a cross-sectional and prospective study in women.” Environ Res 121: 104–109.

7. Beck, R., M. Chandi, M. Kanke, M. Styblo and P. Sethupathy (2019). “Arsenic is more potent than cadmium or manganese in disrupting the INS-1 beta cell microRNA landscape.” Arch Toxicol 93(11): 3099–3109.

8. Bell, R. R., J. L. Early, V. K. Nonavinakere and Z. Mallory (1990). “Effect of cadmium on blood glucose level in the rat.” Toxicol Lett 54(2-3): 199–205.

9. Cai, Y., C. P. Kirschke and L. Huang (2018). “SLC30A family expression in the pancreatic islets of humans and mice: cellular localization in the beta-cells.” J Mol Histol 49(2): 133–145.

10. Chang, K. C., C. C. Hsu, S. H. Liu, C. C. Su, C. C. Yen, M. J. Lee, K. L. Chen, T. J. Ho, D. Z. Hung, C. C. Wu, T. H. Lu, Y. C. Su, Y. W. Chen and C. F. Huang (2013). “Cadmium induces apoptosis in pancreatic beta-cells through a mitochondria-dependent pathway: the role of oxidative stress-mediated c-Jun N-terminal kinase activation.” PLoS One 8(2): e54374.

11. Chimienti, F., A. Favier and M. Seve (2005). “ZnT-8, a pancreatic beta-cell-specific zinc transporter.” Biometals 18(4): 313–317.

12. Coffey, R. and M. D. Knutson (2017). “The plasma membrane metal-ion transporter ZIP14 contributes to nontransferrin-bound iron uptake by human beta-cells.” Am J Physiol Cell Physiol 312(2): C169–C175.

13. Dalmieda, J. and P. Kruse (2019). “Metal Cation Detection in Drinking Water.” Sensors 19(23): 5134–5178.

14. Dalton, T. P., L. He, B. Wang, M. L. Miller, L. Jin, K. F. Stringer, X. Chang, C. S. Baxter and D. W. Nebert (2005). “Identification of mouse SLC39A8 as the transporter responsible for cadmium-induced toxicity in the testis.” Proc Natl Acad Sci U S A 102(9): 3401–3406.

15. El-Muayed, M., M. R. Raja, X. Zhang, K. W. MacRenaris, S. Bhatt, X. Chen, M. Urbanek, T. V. O’Halloran and W. L. Lowe Jr (2012). “Accumulation of Cadmium in Insulin-producing beta-cells.” Islets 4(6): 405–416.

16. El Muayed, M., M. R. Raja, X. Zhang, K. W. MacRenaris, S. Bhatt, X. Chen, M. Urbanek, T. V. O’Halloran and W. L. Lowe, Jr. (2012). “Accumulation of cadmium in insulin-producing beta cells.” Islets 4(6): 405–416.

17. Elinder, C. G., B. Lind, T. Kjellstrom, L. Linnman and L. Friberg (1976). “Cadmium in kidney cortex, liver, and pancreas from Swedish autopsies. Estimation of biological half time in kidney cortex, considering calorie intake and smoking habits.” Arch Environ Health 31(6): 292–302.

18. Fernandez-Twinn, D. S., L. Hjort, B. Novakovic, S. E. Ozanne and R. Saffery (2019). “Intrauterine programming of obesity and type 2 diabetes.” Diabetologia 62(10): 1789–1801.

19. Fitzgerald, R., A. Olsen, J. Nguyen, W. P. Wong, M. El-Muayed and J. Edwards (2020). “Pancreatic Islets Accumulate Cadmium in a Rodent Model of Cadmium-Induced Hyperglycemia.” International Journal of Molecular Sciences 22(1): 360.

20. Friberg, L. (1984). “Cadmium and the kidney.” Environ Health Perspect 54: 1–11.

21. Friedman, R. (2014). “Structural and computational insights into the versatility of cadmium binding to proteins.” Dalton Trans 43(7): 2878-2887.

22. Garrick, M. D., K. G. Dolan, C. Horbinski, A. J. Ghio, D. Higgins, M. Porubcin, E. G. Moore, L. N. Hainsworth, J. N. Umbreit, M. E. Conrad, L. Feng, A. Lis, J. A. Roth, S. Singleton and L. M. Garrick (2003). “DMT1: a mammalian transporter for multiple metals.” Biometals 16(1): 41–54.

23. Girijashanker, K., L. He, M. Soleimani, J. M. Reed, H. Li, Z. Liu, B. Wang, T. P. Dalton and D. W. Nebert (2008). “Slc39a14 gene encodes ZIP14, a metal/bicarbonate symporter: similarities to the ZIP8 transporter.” Mol Pharmacol 73(5): 1413–1423.

24. Hansen, J. B., M. F. Tonnesen, A. N. Madsen, P. H. Hagedorn, J. Friberg, L. G. Grunnet, R. S. Heller, A. O. Nielsen, J. Storling, L. Baeyens, L. Anker-Kitai, K. Qvortrup, L. Bouwens, S. Efrat, M. Aalund, N. C. Andrews, N. Billestrup, A. E. Karlsen, B. Holst, F. Pociot and T. Mandrup-Poulsen (2012). “Divalent metal transporter 1 regulates iron-mediated ROS and pancreatic beta cell fate in response to cytokines.” Cell Metab 16(4): 449–461.

25. Himeno, S., T. Yanagiya and H. Fujishiro (2009). “The role of zinc transporters in cadmium and manganese transport in mammalian cells.” Biochimie 91(10): 1218–1222.

26. Hong, H., Y. Xu, J. Xu, J. Zhang, Y. Xi, H. Pi, L. Yang, Z. Yu, Q. Wu, Z. Meng, W. S. Ruan, Y. Ren, S. Xu, Y. Q. Lu and Z. Zhou (2021). “Cadmium exposure impairs pancreatic beta-cell function and exaggerates diabetes by disrupting lipid metabolism.” Environ Int 149: 106406.

27. Huang, C. C., C. Y. Kuo, C. Y. Yang, J. M. Liu, R. J. Hsu, K. I. Lee, C. C. Su, C. C. Wu, C. T. Lin, S. H. Liu and C. F. Huang (2019). “Cadmium exposure induces pancreatic beta-cell death via a Ca(2+)-triggered JNK/CHOP-related apoptotic signaling pathway.” Toxicology 425: 152252.

28. Jarup, L. and A. Akesson (2009). “Current status of cadmium as an environmental health problem.” Toxicol Appl Pharmacol 238(3): 201–208.

29. Jarup, L., A. Rogenfelt, C. G. Elinder, K. Nogawa and T. Kjellstrom (1983). “Biological half-time of cadmium in the blood of workers after cessation of exposure.” Scand J Work Environ Health 9(4): 327–331.

30. Jin, T., G. F. Nordberg and M. Nordberg (1986). “Uptake of cadmium in isolated kidney cells--influence of binding form and in vivo pretreatment.” J Appl Toxicol 6(6): 397–400.

31. Jolly, Y. N., A. Islam and S. Akbar (2013). “Transfer of Metals from Soil to Vegetables and Possible Health Risk Assessment.” SpringerPlus 15(1): 385–393.

32. Kambe, T., H. Narita, Y. Yamaguchi-Iwai, J. Hirose, T. Amano, N. Sugiura, R. Sasaki, K. Mori, T. Iwanaga and M. Nagao (2002). “Cloning and characterization of a novel mammalian zinc transporter, zinc transporter 5, abundantly expressed in pancreatic beta cells.” J Biol Chem 277(21): 19049–19055.

33. Kolb, H. and S. Martin (2017). “Environmental/Lifestyle Factors in the Pathogenesis and Prevention of Type 2 Diabetes.” BMC Medicine 15(1): 131–142.

34. Kuo, C. C., K. A. Moon, S. L. Wang, E. Silbergeld and A. Navas-Acien (2017). “The Association of Arsenic Metabolism with Cancer, Cardiovascular Disease, and Diabetes: A Systematic Review of the Epidemiological Evidence.” Environ Health Perspect 125(8): 087001.

35. Lawlor, N., J. George, M. Bolisetty, R. Kursawe, L. Sun, V. Sivakamasundari, I. Kycia, P. Robson and M. L. Stitzel (2017). “Single-cell transcriptomes identify human islet cell signatures and reveal cell-type-specific expression changes in type 2 diabetes.” Genome Res 27(2): 208–222.

36. Lawson, R., W. Maret and C. Hogstrand (2017). “Expression of the ZIP/SLC39A transporters in beta-cells: a systematic review and integration of multiple datasets.” BMC Genomics 18(1): 719.

37. Lemaire, K., F. Chimienti and F. Schuit (2012). “Zinc transporters and their role in the pancreatic beta-cell.” J Diabetes Investig 3(3): 202–211.

38. Li, X., M. Li, J. Xu, X. Zhang, W. Xiao and Z. Zhang (2019). “Decreased Insulin Secretion but Unchanged Glucose Homeostasis in Cadmium-Exposed Male C57BL/6 Mice.” J Toxicol 2019: 8121834.

39. Li, Y. Y., C. Douillet, M. Huang, R. Beck, S. J. Sumner and M. Styblo (2020). “Exposure to inorganic arsenic and its methylated metabolites alters metabolomics profiles in INS-1 832/13 insulinoma cells and isolated pancreatic islets.” Arch Toxicol 94(6): 1955–1972.

40. Lortz, S., S. Schroter, V. Stuckemann, I. Mehmeti and S. Lenzen (2014). “Influence of cytokines on Dmt1 iron transporter and ferritin expression in insulin-secreting cells.” J Mol Endocrinol 52(3): 301–310.

41. Martin, E., C. Gonzalez-Horta, J. Rager, K. A. Bailey, B. Sanchez-Ramirez, L. Ballinas-Casarrubias, M. C. Ishida, D. S. Gutierrez-Torres, R. Hernandez Ceron, D. Viniegra Morales, F. A. Baeza Terrazas, R. J. Saunders, Z. Drobna, M. A. Mendez, J. B. Buse, D. Loomis, W. Jia, G. G. Garcia-Vargas, L. M. Del Razo, M. Styblo and R. Fry (2015). “Metabolomic characteristics of arsenic-associated diabetes in a prospective cohort in Chihuahua, Mexico.” Toxicol Sci 144(2): 338–346.

42. Menke, A., E. Guallar and C. C. Cowie (2016). “Metals in Urine and Diabetes in U.S. Adults.” Diabetes 65(1): 164–171.

43. Miyazaki, J., K. Araki, E. Yamato, H. Ikegami, T. Asano, Y. Shibasaki, Y. Oka and K. Yamamura (1990). “Establishment of a pancreatic beta cell line that retains glucose-inducible insulin secretion: special reference to expression of glucose transporter isoforms.” Endocrinology 127(1): 126–132.

44. Mohanasundaram, D., C. Drogemuller, J. Brealey, C. F. Jessup, C. Milner, C. Murgia, C. J. Lang, A. Milton, P. D. Zalewski, G. R. Russ and P. T. Coates (2011). “Ultrastructural analysis, zinc transporters, glucose transporters and hormones expression in New world primate (Callithrix jacchus) and human pancreatic islets.” Gen Comp Endocrinol 174(2): 71–79.

45. Moon, S. S. (2013). “Association of lead, mercury and cadmium with diabetes in the Korean population: the Korea National Health and Nutrition Examination Survey (KNHANES) 2009-2010.” Diabet Med 30(4): e143–148.

46. Nie, X., N. Wang, Y. Chen, C. Chen, B. Han, C. Zhu, Y. Chen, F. Xia, Z. Cang, M. Lu, Y. Meng, B. Jiang, D. J. M and Y. Lu (2016). “Blood cadmium in Chinese adults and its relationships with diabetes and obesity.” Environ Sci Pollut Res Int 23(18): 18714–18723.

47. Nordberg, G. F. (2004). “Cadmium and health in the 21st century--historical remarks and trends for the future.” Biometals 17(5): 485–489.

48. Ohana, E., I. Sekler, T. Kaisman, N. Kahn, J. Cove, W. F. Silverman, A. Amsterdam and M. Hershfinkel (2006). “Silencing of ZnT-1 expression enhances heavy metal influx and toxicity.” J Mol Med (Berl) 84(9): 753–763.

49. Petersmann, A., D. Muller-Wieland, U. A. Muller, R. Landgraf, M. Nauck, G. Freckmann, L. Heinemann and E. Schleicher (2019). “Definition, Classification and Diagnosis of Diabetes Mellitus.” Exp Clin Endocrinol Diabetes 127(S 01): S1-S7.

50. Scherer, G. and H. Barkemeyer (1983). “Cadmium Concentrations in Tobacco and Tobacco Smoke.” Ecotoxicology and Environmental Safety 7(1): 71–78.

51. Schilderman, P. A., E. J. Moonen, P. Kempkers and J. C. Kleinjans (1997). “Bioavailability of soil-adsorbed cadmium in orally exposed male rats.” Environ Health Perspect 105(2): 234–238.

52. Schwartz, G. G., D. II’yasova and A. Ivanova (2003). “Urinary Cadmium, Impaired Fasting Glucose and Diabetes in the NHANES III.” Diabetes Care 26(2): 468–470.

53. Tallkvist, J., C. L. Bowlus and B. Lonnerdal (2001). “DMT1 gene expression and cadmium absorption in human absorptive enterocytes.” Toxicol Lett 122(2): 171–177.

54. Todd, J. N., S. Srinivasan and T. I. Pollin (2018). “Advances in the Genetics of Youth-Onset Type 2 Diabetes.” Curr Diab Rep 18(8): 57.

55. Trevino, S., M. P. Waalkes, J. A. Flores Hernandez, B. A. Leon-Chavez, P. Aguilar-Alonso and E. Brambila (2015). “Chronic cadmium exposure in rats produces pancreatic impairment and insulin resistance in multiple peripheral tissues.” Arch Biochem Biophys 583: 27–35.

56. Wallia, A., N. B. Allen, S. Badon and M. El Muayed (2014). “Association between urinary cadmium levels and prediabetes in the NHANES 2005-2010 population.” Int J Hyg Environ Health 217(8): 854–860.

57. Winslow, J. W. W., K. H. Limesand and N. Zhao (2020). “The Functions of ZIP8, ZIP14, and ZnT10 in the Regulation of Systemic Manganese Homeostasis.” Int J Mol Sci 21(9).

58. Wong, W. P., N. B. Allen, M. S. Meyers, E. O. Link, X. Zhang, K. W. MacRenaris and M. El Muayed (2017). “Exploring the Association Between Demographics, SLC30A8 Genotype, and Human Islet Content of Zinc, Cadmium, Copper, Iron, Manganese and Nickel.” Sci Rep 7(1): 473.

59. Wong, W. P. S., J. C. Wang, M. J. Schipma, X. Zhang, J. R. Edwards and M. El Muayed (2021). “Cadmium-mediated pancreatic islet transcriptome changes in mice and cultured mouse islets.” Toxicol Appl Pharmacol 433: 115756.

